# Reactive Pericytes Lead to Microvascular Dysfunction and Cortical Neurodegeneration During Experimental Autoimmune Encephalomyelitis

**DOI:** 10.1101/2025.02.17.638582

**Authors:** Mohammadreza Yousefi, Ayşe Özkan, Kaan Kutay Özmen, Yiğit Uysallı, Mina Mamipour, Nazan Akkaya, Amir-Reza Dalal Amandi, Abdulsamet Atbaşi, Morteza Heidarzadeh, Şefik Evren Erdener, Naoto Kawakami, Alper Kiraz, Yasemin Gürsoy Özdemir, Atay Vural

**Affiliations:** Koç University Research Center for Translational Medicine (KUTTAM), İstanbul, Türkiye; Koç University Graduate School of Health Sciences, İstanbul, Türkiye; Koç University School of Medicine, Department of Neurology, İstanbul, Türkiye; Koç University, Department of Physics, İstanbul, Türkiye; Koç University, Department of Electrical and Electronics Engineering, İstanbul, Türkiye; İzmir Bakırçay University, Department of Physiology, İzmir, Türkiye; University of Tabriz, Department of Animal Biology, Faculty of Natural Science, Tabriz, Iran; Institute of Neurological Sciences and Psychiatry, Hacettepe University, Ankara, Türkiye; Institute of Clinical Neuroimmunology, University Hospital, Ludwig-Maximilians-Universität München, Munich, Germany

**Keywords:** Pericytes, cerebrovascular reactivity, neurovascular unit, experimental autoimmune encephalitis, multiple sclerosis, neurodegeneration

## Abstract

The mechanisms underlying neurodegeneration in multiple sclerosis remain incompletely understood. In this study, we aimed to investigate the role of vascular dysfunction in cortical neurodegeneration using a chronic cranial window model of experimental autoimmune encephalomyelitis in mice. After the induction of experimental autoimmune encephalomyelitis with myelin oligodendrocyte glycoprotein peptides in C57BL/6J mice, we assessed cerebrovascular reactivity though a chronic cranial window using laser speckle contrast imaging and intrinsic optical signal imaging in awake animals. We observed a significant reduction in cortical cerebrovascular reactivity during peak inflammation in the EAE group, as detected by laser speckle contrast imaging after 5% hypercapnia (*p=*0.04) and optical signal imaging after whisker stimulation (*p=*0.008). Histological analysis revealed a diffuse increase in CD13+ pericyte coverage (*p=*0.001), accompanied by focal IgG deposition within the microvascular lumen (*p=*0.04) and increased amount of CD45+ leukocytes stalled in microvessels (*p=*0.03) in the cortex of experimental autoimmune encephalomyelitis mice. Microglial activation was also present in the cortex of experimental autoimmune encephalomyelitis mice (*p*=0.04) and was particularly evident around microvessels with IgG deposition. Subpial and intracortical foci exhibiting loss of NeuN reactivity (*p=*0.03) and axonal loss (*p=*0.007) were detected in experimental autoimmune encephalomyelitis, but not in control mice. Altogether, these results demonstrate that microvascular function and neurovascular unit elements are globally affected in the cortex during autoimmune neuroinflammation and is related to neurodegeneration.

## Introduction

Neurodegeneration is evident from the earliest stages of multiple sclerosis (MS) (1, 2). While the immune mechanisms driving neuroinflammation are well understood, the processes linked to neurodegeneration remain poorly characterized, and effective therapies targeting neurodegeneration in MS are still lacking. Several mechanisms have been proposed to contribute to neurodegeneration in MS, including hypoxia resulting from mitochondrial dysfunction (3, 4). Another potential explanation is the “vascular hypothesis,” which suggests that dysfunction of the blood-brain barrier (BBB) and/or the neurovascular unit (NVU) may contribute to neurodegeneration. This hypothesis has two key components: the first is based on observations of BBB leakage and the neurotoxic effects of serum proteins such as fibrinogen; the second component suggests that cerebrovascular function is disrupted in MS, a theory supported by studies in patients with MS (pwMS) (5, 6).

The NVU regulates cerebral blood flow (CBF) in response to neuronal activation by dilating cerebral microvessels and increasing local perfusion in a process called neurovascular coupling (NVC) (7, 8). The NVU consists of endothelial cells, pericytes that ensheath microvessels, astrocytes, and neurons. NVU dysfunction refers to the breakdown of coordination between neurons and blood vessels in the central nervous system (9, 10). Dysfunction of the NVU has been identified as an early event and a significant contributor to the pathogenesis of Alzheimer’s disease and other neurodegenerative disorders (11). In pwMS, several studies have examined NVU function and cerebrovascular reactivity (CVR) (12). Some studies on CVR have shown reduced CVR in pwMS compared to healthy controls (HC), while others found no significant differences, likely due to variability in study cohorts. Similarly, research on NVC has produced mixed results, with some studies indicating decreased NVC in pwMS, while others have found no significant changes (12). This inconsistency highlights the need for animal model studies to further explore the role of NVC in MS pathogenesis.

Pericytes play a crucial role in regulating cerebral blood flow, and their loss leads to impaired vascular reactivity and NVC (13). The relationship between pericyte loss, NVU dysfunction, and neurodegeneration has been demonstrated in Alzheimer’s disease, ALS, Huntington’s disease, and other neurodegenerative conditions (11, 14, 15). However, the contribution of pericytes to MS pathogenesis, particularly in relation to cerebrovascular dysfunction and neurodegeneration, remains unclear.

In this study, we used the myelin oligodendrocyte glycoprotein (MOG)-induced experimental autoimmune encephalomyelitis (EAE) model to investigate the contribution of NVC, CVR, and pericytes to the neurodegeneration observed in pwMS. The MOG-induced EAE model is particularly suitable for studying the mechanisms underlying diffuse cortical neurodegeneration, which occurs independently of the acute lesions formed by leukocyte infiltration in pwMS. In this model, immune cell infiltration and lesion formation predominantly occur in the spinal cord, making it an appropriate model for examining the remote effects of cerebrospinal fluid (CSF) soluble factors (cytokines, reactive oxygen species, etc.) released during neuroinflammation on the cortex. Our findings demonstrate that NVU function is disrupted in the cortex, which is associated with an increase in cortical pericyte coverage, and the upregulation of genes related to cytoskeleton organization and contraction. These changes were accompanied by microglial activation, a loss of NeuN reactivity, and neurite loss, which predominantly occurred around affected microvessels.

## Material and Methods

### Animals

Animal experiments were conducted at the Koç University Animal Research Laboratory in accordance with the ARRIVE guidelines and regulations approved by the Institutional Animal Care and Use Committee of Koç University (approval no: 2019.HADYEK.021). All animals were individually housed in standard cages under controlled conditions, including temperature, humidity, and a 12-hour light-dark cycle, with unrestricted access to food and water. Male wild type C57BL/6J mice, aged 8-12 weeks and weighing 20-25 g, were used for the experiments.

### Cranial Window Surgery

Closed cranial windows were prepared for longitudinal imaging of CBF by using the methodology of Kilic et al. (16). Preoperative analgesia was provided with intraperitoneal injections of Buprenorphine (0.05 mg/kg, diluted in saline) and Dexamethasone (4.8 mg/kg) administered 6 and 2 hours before surgery, respectively, to alleviate pain and minimize brain swelling. Anesthesia was induced with 3% isoflurane, followed by an intraperitoneal injection of 250 μl of Ketamine/Xylazine solution (100/10 mg/kg). The anesthetized mouse was then secured in a sterile stereotaxic frame on a heated pad. A 5-mm circular craniotomy was performed in the left hemisphere, targeting the barrel field of the somatosensory cortex, centered 1 mm posterior and 1.5 mm lateral to the bregma. To mitigate potential bleeding and prevent skull regrowth under the window, the cortical surface was carefully cleaned of blood, and sterile room-temperature artificial cerebrospinal fluid (ACSF) was applied. A custom-made T-shaped glass plug (Supplementary Method) constructed from one 5-mm round coverslip and three 3-mm round coverslip was placed over the exposed cortex. The plug was secured using a thin layer of cyanoacrylate adhesive (502 Super Glue) applied around the edge, ensuring no glue reached the brain tissue. Next, a rectangle-shaped 3D-printed polylactic acid head plate (Supplementary Method) was affixed to the skull using dental cement, leaving the cranial window exposed to facilitate head fixation during imaging. Postoperative care was provided by administering a cocktail of Sulfamethoxazole (1 mg/ml), Trimethoprim (0.2 mg/ml), and Ibuprofen (1 mg/ml) in drinking water for at least 10 days to reduce inflammation (16). To acclimate the mice to head-fixation during awake imaging, they were trained daily for over a period of 7-10 days, ensuring they could be maintained in the head plate restraint comfortably without excess movement during *in vivo* imaging.

### Training Awake Mice for Acclimation to Head Plate Restraint and Sensory Stimulation

Mice were trained and habituated to the custom-made head plate and its restraint through a brief, structured training procedure. Before acclimation, the mice participated in daily handling sessions in the experimental room, with consistent light, noise space and instrument, for at least 7-10 days, with each session lasting 5 minutes at the beginning of the day. This continued until the mice revealed relaxed behavior, allowing them to freely explore the experimenter’s hand and arm. Once this relaxed behavior was observed, the mice were placed into the transparent tube of the head plate restraint for 5 minutes and then fixed to the restraint. The mice were trained for up to 30 minutes, with the duration gradually increasing each day (5, 10, 15, 20, 25, and finally 30 minutes). As the acclimation progressed, the duration of holding the mice’s heads gradually increased, and they firmly secured without showing any movement to the head plate restraint. To promote acclimation and reduce stress of stimulations (air puff whisker pad and hypercapnia), accompanied with the animal habituation to the restraint, mice were exposed to air puff and %5 CO_2_ for one minute. Ultimately, mice were trained and acclimated to the head plate restraint and its transparent tube, along with the stimulation methods, for 30 minutes each day from day 7-10.

### Whisker and Hypercapnia Stimulations

Cerebrovascular reactivity was assessed using two distinct stimulation protocols. First, whisker stimulation was conducted using a custom device that delivered standardized air puffs. The protocol involved 20-second air puff delivery with a pulse duration of 50 milliseconds at 3 Hz, repeated ten times with 40-second intervals between stimuli. Second, hypercapnia stimulation was performed by ventilating the mice with 5% CO2 through a mask. The hypercapnia protocol consisted of a 60-second exposure to 5% CO2, followed by 60-second intervals, repeated five times. Each whisker and hypercapnia stimulation cycle were composed of non-stimulation, stimulation, and relaxation phases (Supplementary Fig. 1A, B).

### Laser Speckle Contrast Imaging and Data Analysis

Laser Speckle Contrast Imaging (LSCI) (Perimed, Järfälla, Sweden) was used to assess cerebral blood flow (CBF) responses to whisker stimulation and 5% hypercapnia in awake mice. The head of the mice was fixed using a head plate holder and the LSCI camera was positioned 10 cm above the cranial window for imaging. Scanning was performed at 10 frames per second over a 5×5 mm field to track speckle patterns within the illuminated region, using the Pimsoft software (Perimed, Järfälla, Sweden).

For data analysis, regions of interest (ROIs) were manually defined within the exposed cortex under the cranial window, selecting three identical spherical 10×10-pixel regions beyond the rim or slightly farther from large vessels (0.3–0.8 mm away from arterioles) (Supplementary Fig. 1A). For each round of the ten stimulation sessions, a 10-second period from each session was selected, excluding any signal drift or changes (Supplementary Fig. 1B). This step is essential to account for minor movements in the brain and cerebral arteries caused by breathing and heartbeat during recording. Within the interaction of each time of interest (TOI) and the defined ROIs, the mean blood flow index was calculated and normalized to the baseline signal. Finally, the average normalized response of the perfusion signal to stimulation across all sessions was calculated.

### Intrinsic Optical Signal Imaging and Data Analysis

Intrinsic Optical Signal Imaging (IOSI) is an optical technique used to monitor in vivo blood flow changes by detecting alterations in hemoglobin content in response to brain activity, utilizing diffuse light (17, 18). Oxyhemoglobin and deoxyhemoglobin have similar absorption values at common wavelengths, but changes in hemoglobin absorption can be linked to variations in factors such as blood flow and metabolic rates. At 530 nm and other green light wavelengths, the molar extinction coefficients of oxyhemoglobin and deoxyhemoglobin are nearly identical, facilitating the detection of CBF and total hemoglobin changes induced by brain stimulation(19–22).

In this study, IOSI was performed using a custom-made setup (Supplementary Material) under a 570 nm wavelength LED (green light source) (Supplementary Fig. 1C). Images were acquired using MATLAB (R2021b) at 5 frames per second, for 10 minutes at a resolution of 256×256 pixels.

For data analysis, three ROIs (10×10 pixels in size) were defined, preferably in regions showing peak signal changes and positioned away from large vessels (Supplementary Fig. 1C). Then, a 4 mm window segmentation was applied to avoid any interference from surrounding noise during signal reception. A Gaussian kernel spatial filter was then applied to the segmented data to obtain binary data (Supplementary Fig. 1D). After defining the ROIs, 10-second intervals from each stimulation session were selected, excluding any signal drift or changes. The perfusion signal for each stimulation was then normalized to zero, serving as a baseline perfusion cut-off for each pixel in the ROI. This step also allowed for re-alignment of the recorded frames and eliminated any initial positive or negative signal drift, which is essential as breathing and heartbeat can cause slight movement in the brain and cerebral arteries during recording. Finally, the average signals in response to stimulation were calculated using customized MATLAB code.

### Induction and Scoring of EAE

To induce EAE, mice were anesthetized with 2% isoflurane in 2 L/min oxygen and then subcutaneously injected with 0.1 ml of myelin oligodendrocyte glycoprotein (MOG35-55) peptide emulsified in complete Freund’s adjuvant (CFA) (Hooke Laboratories, EL-2110) into the upper and lower back regions. On days 1 and 2 post-immunization, 135 ng of pertussis toxin were administered intraperitoneally (23, 24). Of note, we adjusted the dose of pertussis toxin to 35% higher than the average amount recommended in the kit’s manual, to guarantee prominent inflammation in the EAE cohort.

During the first week after immunization, the animals were housed in a calm, noise-free environment to minimize stress, which could potentially affect the outcomes of EAE. Starting from the second week, weight and clinical scores were monitored every other day. Clinical scores were recorded in a blinded manner, following the EAE scoring guidelines provided by the manufacturer: 0 = no clinical signs; 0.5 = flaccidity of the tail tip; 1 = completely limp tail; 1.5 = limp tail with unilateral hind leg weakness; 2 = limp tail with bilateral hind leg weakness; 2.5 = limp tail with bilateral hind leg dragging; 3 = limp tail with complete bilateral hind leg paralysis; 3.5 = limp tail with complete bilateral hind leg paralysis and circling behavior when placed on its side; 4 = limp tail with complete paralysis of both hind legs and partial paralysis of the front legs; 4.5 = complete paralysis of both hind and front legs, with no ability to move; 5 = moribund or deceased.

### Tissue staining

Mice were anesthetized with 2% isoflurane, after which organs were transcardially perfused with ice-cold DPBS, followed by 4% paraformaldehyde (PFA). The brains and spinal cords were then removed and post-fixed in 4% PFA overnight. Following post-fixation, the tissues were transferred to 30% sucrose solution overnight for dehydration and immersion. The organs were subsequently embedded and frozen in the OCT gel (DDK Italia; #22-118). Cryopreserved tissues were sectioned into 20-μm slices using a cryostat (Leica CM1950) and mounted on Superfrost Plus slides. Then, sections were permeabilized with 2% Triton X-100 (v/v, Sigma-Aldrich; X100) for 15 minutes and then blocked using a buffer containing 5% Normal Goat Serum (NGS) (v/v, Cell Signaling; #5425), 2% Bovine Serum Albumin (BSA) (w/v, Sigma-Aldrich; #A9418), and 0.2% Triton X-100 for 60 minutes at room temperature. After blocking, the sections were incubated overnight at 4°C with primary antibodies diluted in the blocking buffer. The following primary antibodies were used: Rat anti-Mouse CD13 (1:100; Bio-Rad; MCA2183), Rabbit anti-Mouse CD105 (1:200; Abcam; ab221675), Rat anti-Mouse CD31 (1:200; BD Pharmingen; 550274), Dylight 488-conjugated tomato-lectin (1:300; ThermoFisher; DL-1174), Rabbit anti-NeuN (1:200; Abcam; ab177487), Rat anti-Mouse CD45 (1:200; BioLegend; 103102), and Rabbit anti-Neurofilament heavy polypeptide (1:200; Abcam; ab207176).

After overnight incubation, sections were washed in DPBS and then incubated with fluorophore-conjugated secondary antibodies for 1 hour at room temperature (Supplementary Table 1). Sections were washed again in DPBS and mounted using a mounting solution (1:1 DPBS and glycerol) containing 10 mg/ml Hoechst (ThermoFisher; Hoechst 33342).

### Imaging and Image Analysis

The Leica DMI8 SP8 confocal laser scanning microscope, equipped with 10X/0.3 NA, 20X/0.75 NA and 40X/1.10 NA water immersion objective and a Hybrid detector, was used to capture images of immunofluorescent tagged slices. Image analysis was carried out by two blinded investigators using the National Institutes of Health ImageJ software (version 1.54f) and an inter-investigator kappa score was calculated (K=0.92). Analysis of CD13 positive pericyte coverage and numbers, neurite density (with anti-SMI-32 staining), NeuN positive neurons, IgG deposition and leakage, and CD45 positive leukocytes were done as described in the Supplementary Methods.

### Culture of Primary Human Brain Vascular Pericytes and Incubation with Hydrogen Peroxide

Human brain vascular pericytes (HBVP; ScienceCell, USA, #1200) were cultured in pericyte medium (ScienceCell, #1201) supplemented with 2% FBS, 1% pericyte growth supplement (ScienceCell, #1252), and 1% penicillin/streptomycin solution. The cells were incubated at 37°C with 5% CO2 until they reached 80-90% confluency in T25 or T75 culture flasks, with the medium being refreshed every 2-3 days. To investigate the effect of hydrogen peroxide (H_2_O_2_) on pericytes, HBVPs were seeded into 6-well plates at a density of 100,000 cells per well. After 24 hours, the cells were treated with 1 mM H_2_O_2_ and then collected for quantitative real-time polymerase chain reaction (qPCR) analysis following 6 hours of incubation.

### Real-Time Polymerase Chain Reaction

RNA was isolated from harvested cell pellets of HBVPs using the RNA Miniprep Plus Kit (Zymo Research, R1055) according to the manufacturer’s instructions. A total of 500 ng of isolated RNA was used to synthesize cDNA via reverse transcription with random primers (Bio-Rad, iScript cDNA Synthesis Kit, Cat. no. 1708891). qPCR analysis of *TPM1, TPM4, MAP3K20, MYLK, GUCY1A1, GUCY1B1, CALM1, MYL6, MYL9* was then performed using the miScript SYBR® Green PCR Kit (Qiagen, 218073) on a LightCycler 480 Instrument (Roche Diagnostics). Primers were designed using the NCBI Primer Designing Tool and checked with the Gene Runner tool (Version 6.5.52) (Supplementary Table 2). The PCR protocol included an initial denaturation at 95°C for 5 minutes, followed by 40 cycles of 95°C for 15 seconds, annealing at primer-specific Tm for 30 seconds, and extension at 72°C for 30 seconds, concluding with a final extension at 72°C for 5 minutes. GAPDH cDNA was amplified as a reference gene, and the critical threshold (CT) value of GAPDH was subtracted from the CT value of the target mRNA to calculate the ΔCT value. The quantity of target mRNA was then determined using the formula 2^(-ΔCT). All experiments were performed in triplicate with three technical replicates each.

### Statistical analysis

Statistical analyses were conducted using GraphPad Prism software (version 10.2.0, San Diego, California, USA). The Mann-Whitney test was used to compare CBF changes in LSCI and IOSI measurement. We compared pericyte coverage variables, pericyte count, NeuN-positive cells, neurotic density, IgG deposition, intravascular CD45+ and extravascular CD45+ cells with unpaired t-test analysis. Differences were considered statistically significant at p < 0.05. Data analyzed with the Mann-Whitney test are presented as the median with interquartile range, while data analyzed with the unpaired t-test are presented as the mean ± SD.

## Results

### Cerebrovascular reactivity and neurovascular coupling are disturbed in the normal appearing cortex during the peak of EAE

To investigate the impact of neuroinflammation in normally appearing cortex, we utilized the MOG peptide induced EAE model (23). In this model, leukocyte infiltration and lesion formation occur in the spinal cord, largely sparing the brain. Therefore, it is a suitable model to investigate the remote effects of soluble factors released during central nervous system (CNS) inflammation in the cortex, in analogy to the normally appearing gray matter (NAGM) in people with MS (pwMS). The timeline of our experimental procedure is shown in Fig. 1A. The clinical score and weight of animals is shown in Fig. 1B and C, respectively.

**Figure 1.**
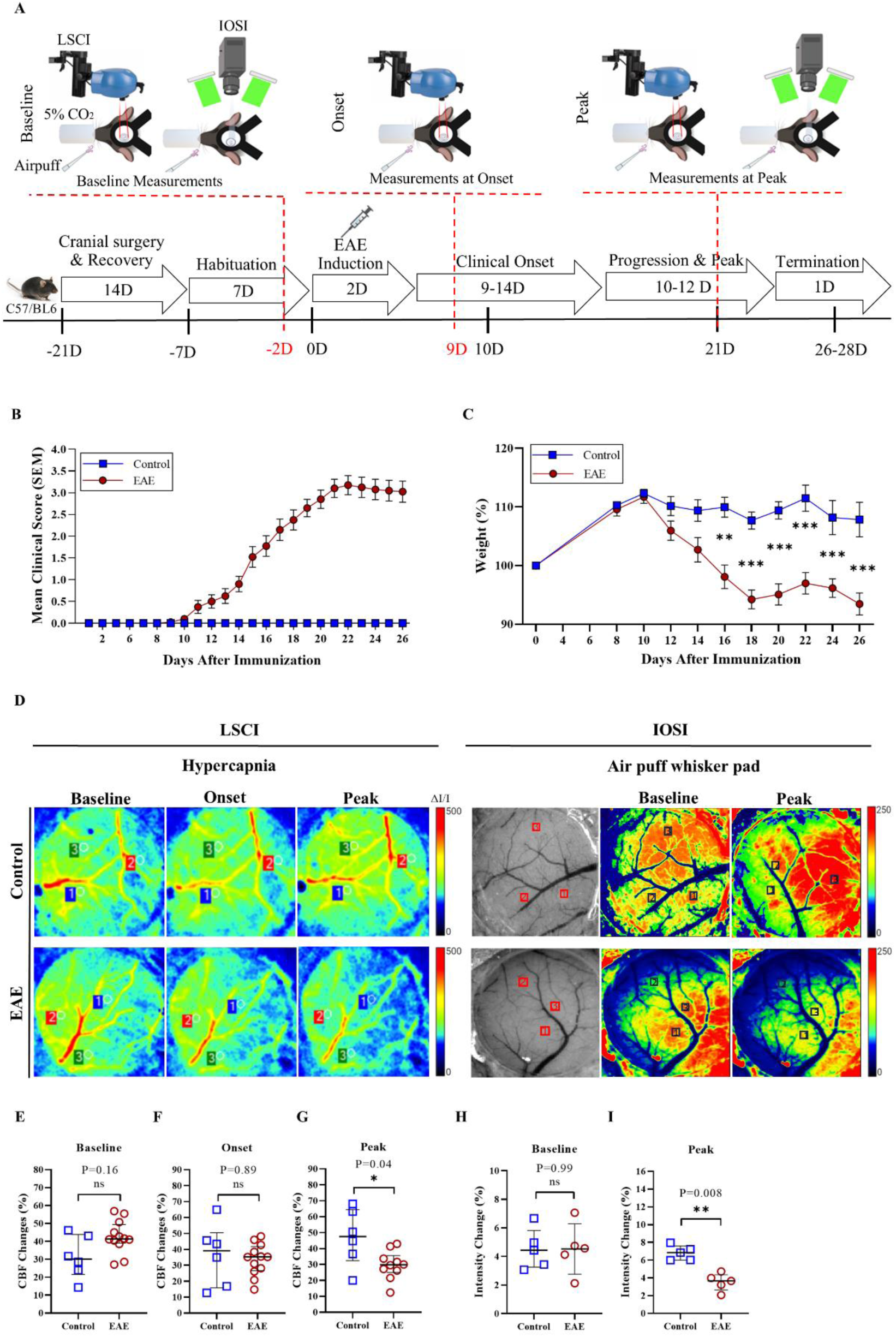
Cerebrovascular reactivity and Neurovascular coupling is disturbed in the cortex during EAE. **A.** The experimental timeline was conducted following cranial window surgery, followed by one week of acclimatization. EAE induction was performed on day 0, after which the clinical score and weight changes were recorded from the onset of symptoms. CBF measurementswere performed at three time points: Baseline, Onset, and Peak respectively. At the final stage of the experiment (days 26-28), animals were transcardially perfused. **B, C.** Clinical symptoms and weight loss in the EAE model were monitored for 28 days. B shows the progression of a typical clinical score, peaking at 3.0 score on day 22, with subsequent stabilization. C presents normalized weight changes, with significant differences between the experimental and control groups from day 14 until the termination day. Statistical analysis was performed using two-way ANOVA with repeated measures and multiple comparisons test (Sidak), with n=6 in the control group and n=20 in the EAE group. P values for days 16, 18, 20, 22, 24, and 26 were **p=0.002, ***p=˂0.001, ***p=˂0.001, ***p=˂0.001, ***p=˂0.001, and ***p=˂0.001, respectively. Error bars represent the mean ± SEM. **D.** Heat maps show CBF changes in control and EAE mice at various time points following hypercapnia and air puff whisker stimulation. Perfusion and intensity changes in 10 whisker and 5 hypercapnia stimulation ROIs interacting with TOIs were calculated and averaged using Pimsoft for LSCI and MATLAB for IOSI. At the peak time points, hypercapnia stimulation in LSCI and air puff whisker stimulation in IOSI resulted in mean perfusion changes in EAE animals. **E-G.** Panels **E** and **F** show no significant differences in CBF changes during the baseline and onset time points following hypercapnia stimulation. In contrast, panel **G** indicates significant CBF changes of up to 37% in EAE mice during the peak time point following hypercapnia stimulation. The P values at the peak time point were *p=0.04. **H, I.** Panels **H** depict quantification results that indicate no significant difference in intensity changes in EAE mice at baseline time points compared to control mice. However, panel **I** indicates a significant decrease of up to 53% in intensity in EAE mice at the peak time point following air puff whisker stimulation. P values for the peak time points were **p= 0.008. The statistical analysis was performed using the Mann-Whitney U test, with n=6 in the control group, n=12 in EAE group for LSCI, and n=5 in both the control and EAE groups for IOSI. Error bars represent median with interquartile range.

We initially assessed cortical microvascular function by measuring CVR with LSCI after hypercapnia induction with 5% CO_2_, and NVC with IOSI after whisker pad stimulation with air puff. We found that the increase in CBF in response to the hypercapnia challenge was similar between the EAE group and controls during baseline measurements and the onset of EAE (Fig. 1D-F). However, CVR exhibited a decrease during the peak of inflammation in the EAE group compared to that in the controls, as evidenced by a 37% lower increase in CBF following hypercapnia induction (Fig. 1D, G; *p*=0.04).

NVC was also disturbed during the peak of EAE. Baseline IOSI recordings after air puff whisker stimulation did not show any differences in cerebral blood volume (CBV) changes between the two groups (Fig. 1D, H). However, a lower increase in CBV was observed in the sensory cortex after whisker stimulation during the peak of inflammation in EAE mice compared to controls (Fig. 1D, I; *p*=0.008).

Overall, these data indicate that both CVR and NVC are compromised during EAE, indicating pericyte dysfunction, which is known as the key regulator of cortical vascular physiology.

### Cortical pericytes react to inflammation by increasing vascular coverage

After documenting disturbed pericyte function, we next assessed pericyte coverage and number. We found that CD13+ pericyte coverage was significantly increased throughout the cortex (103%) and hippocampus (118%) in the EAE group compared to controls (Fig. 2A, B; *p*=0.001, and Supplementary Fig. 2A, B; *p*=0.002, respectively). Pericyte count remained unchanged in both the cortex and hippocampus of EAE-induced mice compared with controls (Fig. 2C, and Supplementary Fig. 2C, respectively). These findings indicate that cortical pericytes sense and react to inflammation even in the absence of direct leukocyte infiltration.

**Figure 2.**
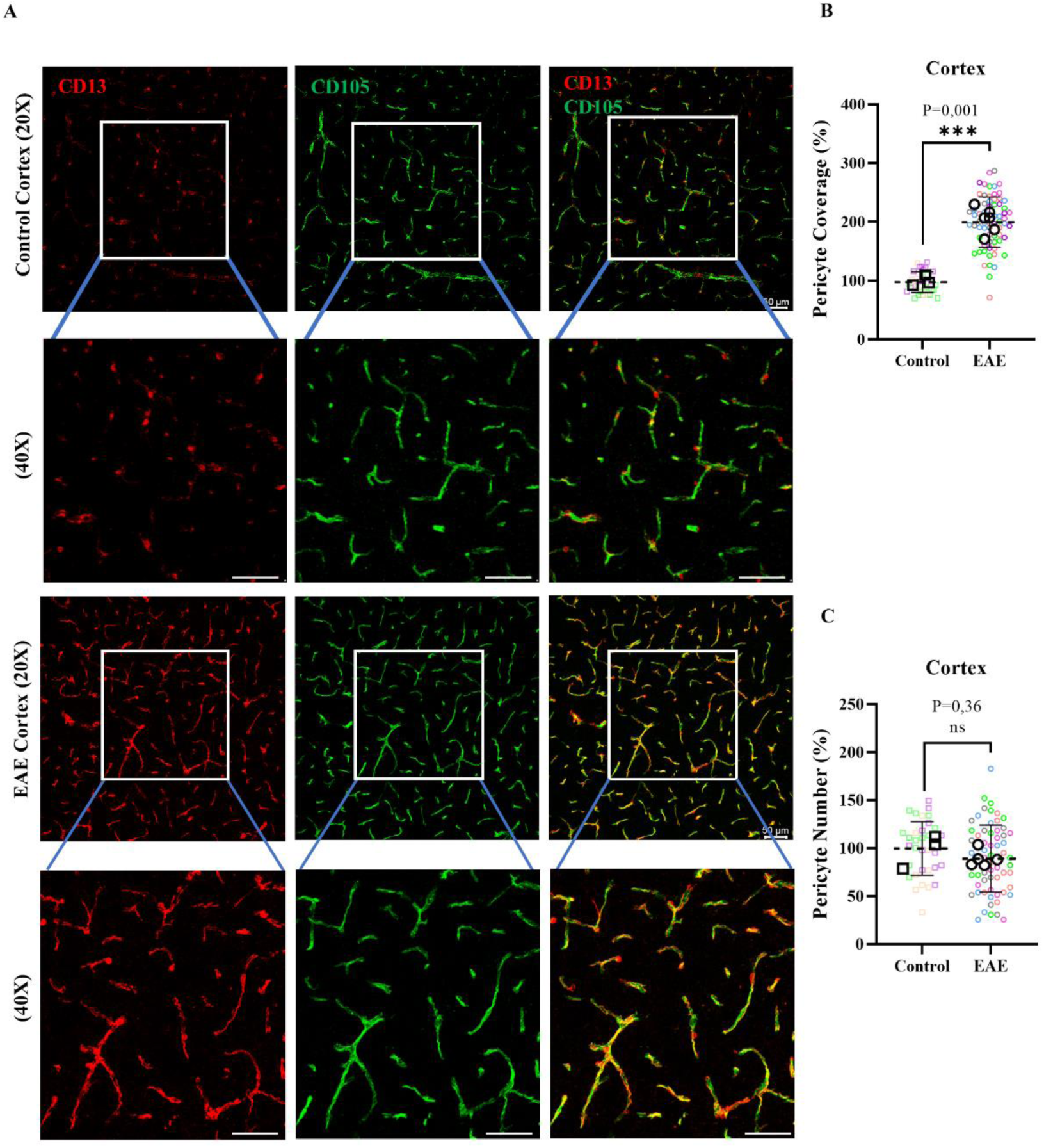
Pericyte coverage is increased in response to inflammation around cortical microvessels during EAE. **A.** Immunofluorescence staining reveals pericyte coverage (CD13, red) and endothelial proliferating cells marked by CD105 (vasculature, green) in the cortex of control and induced EAE mice. Representative confocal images were taken using 20X objectives and magnification was applied with 40X in both control and EAE mice. **B.** Quantification of vascular pericyte coverage reveals a %103 increase in the cortex of induced EAE mice compared to control mice (***p≤0.001). **C.** Quantification of the number of pericyte cell bodies on the microvasculature demonstrates no significant difference in the cortex of induced EAE mice compared to control mice (^ns^ p=0.36). Statistical analysis was performed using a two-tailed unpaired t-test, with n=3 in the control group and n=6 in the EAE group. Error bars represent the mean with +/− SEM. Scale bars in all images represent 50µm.

### Cultured human brain vascular pericytes respond to H2O2 by upregulating cytoskeletal genes

We hypothesized that cortical pericytes increase their vascular coverage in response to soluble factors released remotely by infiltrating leukocytes in the spinal cord lesions formed during the MOG-induced EAE model. To investigate whether human brain vascular pericytes react to reactive oxygen species that are known to be released during neuroinflammation, we measured the expression levels of cytoskeleton- and contraction-related genes in response to H_2_O_2_. After 6 hours of incubation with 1 mM H_2_O_2_, we detected significant upregulation of genes related to cytoskeletal organization and contraction, including *TPM1, TPM4, MAP3K20, MYLK, CALM1, MYL6,* and *MYL9* (Fig. 3). Additionally, the genes *GUCY1A1* and *GUCY1B1*, which encode the two subunits of soluble guanylate cyclase were also upregulated. These findings suggest that the observed increase in pericyte coverage may result from oxidative stress-induced cytoskeletal reorganization under inflammatory conditions.

**Figure 3.**
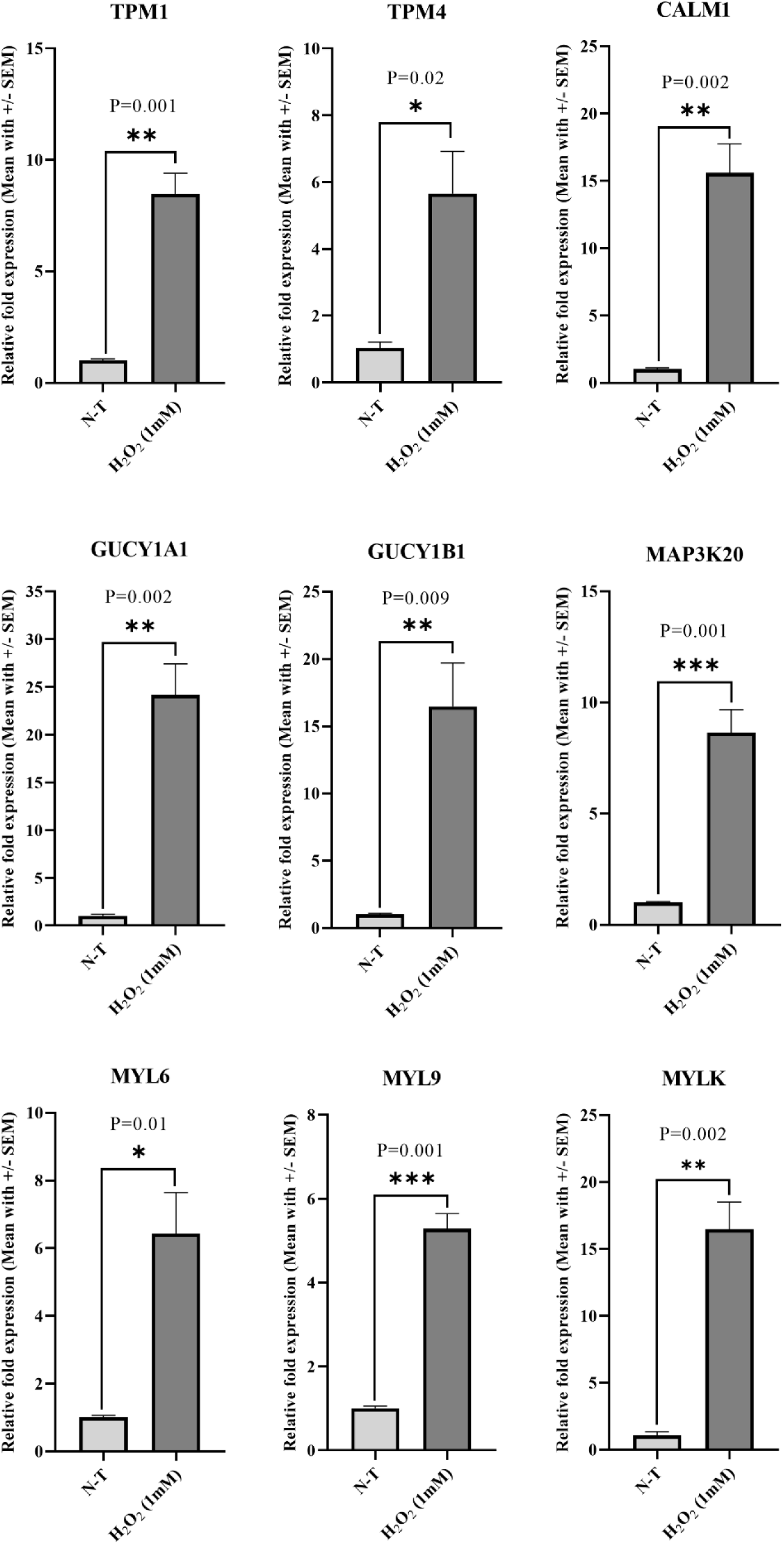
Real-time PCR analysis of cytoskeletal-related genes in HBVP cells following H_2_O_2_ treatment. Relative fold expression analysis revealed that 6-hours exposure to 1mM H_2_O_2_ had the potential to increase the transcription of cytoskeletal-related genes in HBVP cells compare to the None-Treated **(N-T)** group (TPM1; **p=0.001, TPM4; *p=0.02, CALM1; **p=0.002, GUCY1A1; **p=0.002, GUCY1B1; **p=0.009, MAP3K20; ***p=0.001, MYL6; *p=0.01, MYL9; ***p=0.001, MYLK; **p=0.002). The two-tailed unpaired t-test was performed between two groups, and data are the values of three independent experiments. Error bar represented mean with +/− SEM.

### Intraluminal IgG deposition and ICAM1 expression are increased in cortical microvessels during EAE

Pericyte reactivity and dysfunction can lead to BBB disruption and microcirculatory deficits. Therefore, we next assessed whether there is BBB leakage or leukocytic infiltration in the cortex during EAE using anti-IgG and anti-CD45 staining, respectively.

Overall, the IgG signal was significantly increased in both the cortex and hippocampus of EAE-induced mice compared to controls (Fig. 4A, B; *p*=0.04 and Supplementary Fig. 3A, B; p=0.04, respectively). Upon detailed examination of the IgG staining pattern, we observed that IgG deposition was present in focal areas within the cortex (Supplementary Fig. 3C) and predominantly localized intraluminally within microvessels (Fig. 4C). Importantly, pericyte coverage increased more prominently on microvessels with IgG deposition than in microvessels without IgG deposition (Fig. 4C).

**Figure 4.**
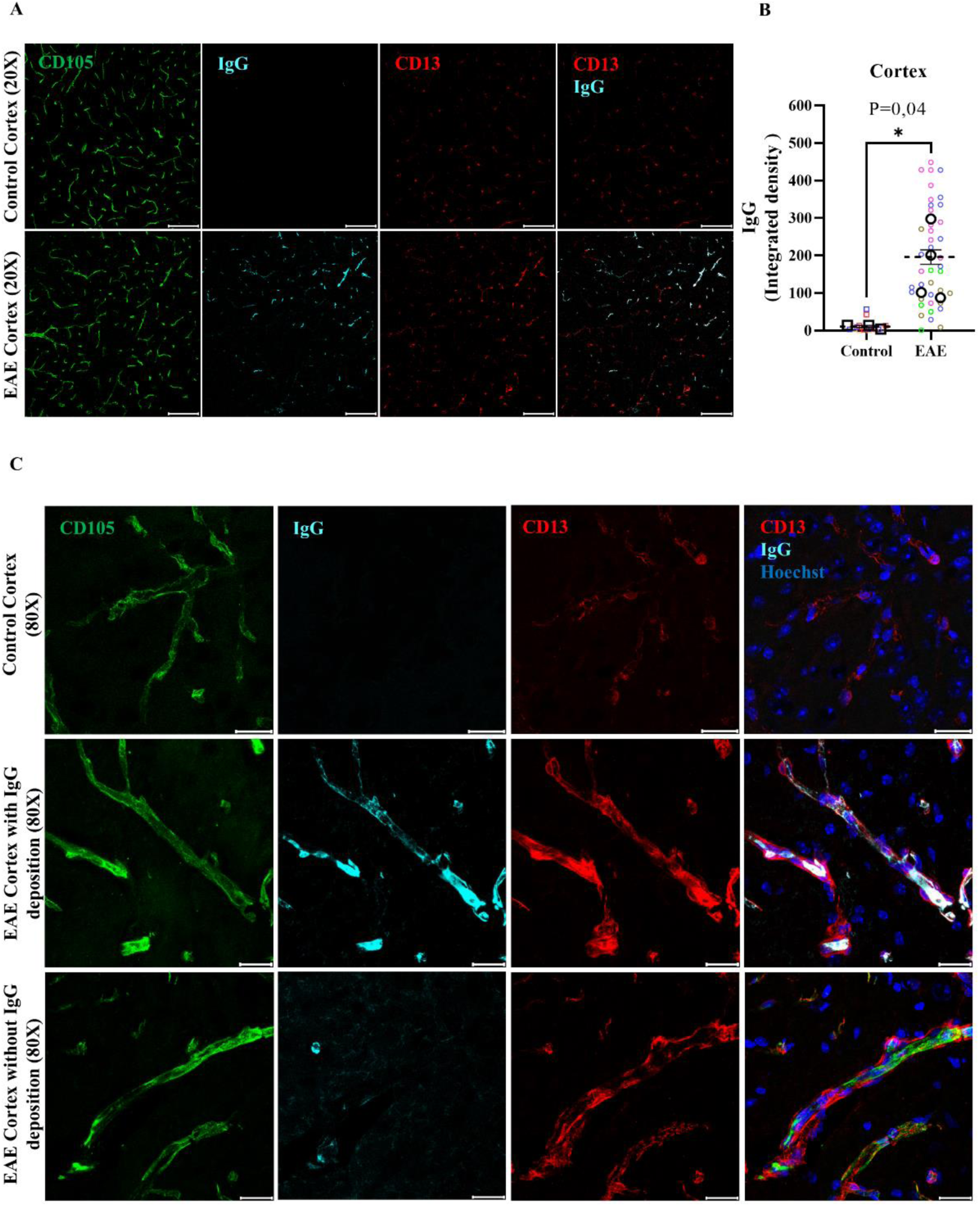
Increased luminal IgG deposition and ICAM1 expression in the cortical microvessels during EAE. **A.** Confocal microscopy images show plasma-derived IgG stained (cyan), along with CD105-stained vessels (green) and CD13-stained pericytes (red), in the cortex of control and induced EAE mice. Images were taken by 20X objectives. **B.** Quantification of plasma-derived IgG in the cortex reveals a substantial ∼ 161% (over 16-fold) increase in luminal IgG deposition in induced EAE mice compared to control mice (*p= 0.04). **C.** Plasma-derived IgG shows significant an increase in the luminal region of the microvasculature, accompanied by increased pericyte coverage in the same area. Statistical analysis was performed using a two-tailed unpaired t-test (n=3 in the control group and n=4 in the EAE group). Error bar represents mean with +/− SEM. Scale bars are 100 µm in A and 10µm in B, respectively.

Next, we assessed ICAM1 expression on cortical microvessels and found increased expression in selected microvessels in the EAE group, whereas there was no detectable ICAM1 expression in control mice (Supplementary Fig. 4).

### Increased leukocyte stalling, but minimal leukocyte extravasation in the EAE cortex

Pericyte dysfunction and endothelial activation, as evidenced by IgG deposition and increased ICAM1 levels, may contribute to enhanced leukocyte stalling and extravasation. Therefore, we quantified the intravascular and extravascular CD45+ leukocytes in the cortex. Our findings revealed a significant increase in the number of CD45+ leukocytes within the intravascular compartment of capillaries in EAE mice compared to controls (Fig. 5A, B; *p*=0.03). A minimal number of extravasated CD45+ leukocytes were also detected in the cortex of EAE mice, whereas none was present in the controls (Fig. 5C; *p*=0.03). Of note, in all stained sections across animals, there were only two microvessels in the cortex with prominent extravascular CD45+ leukocyte infiltration, verifying that the cortex, unlike the spinal cord white matter, is largely devoid of lesion formation characterized by leukocyte extravasation (Supplementary Fig. 5).

**Figure 5.**
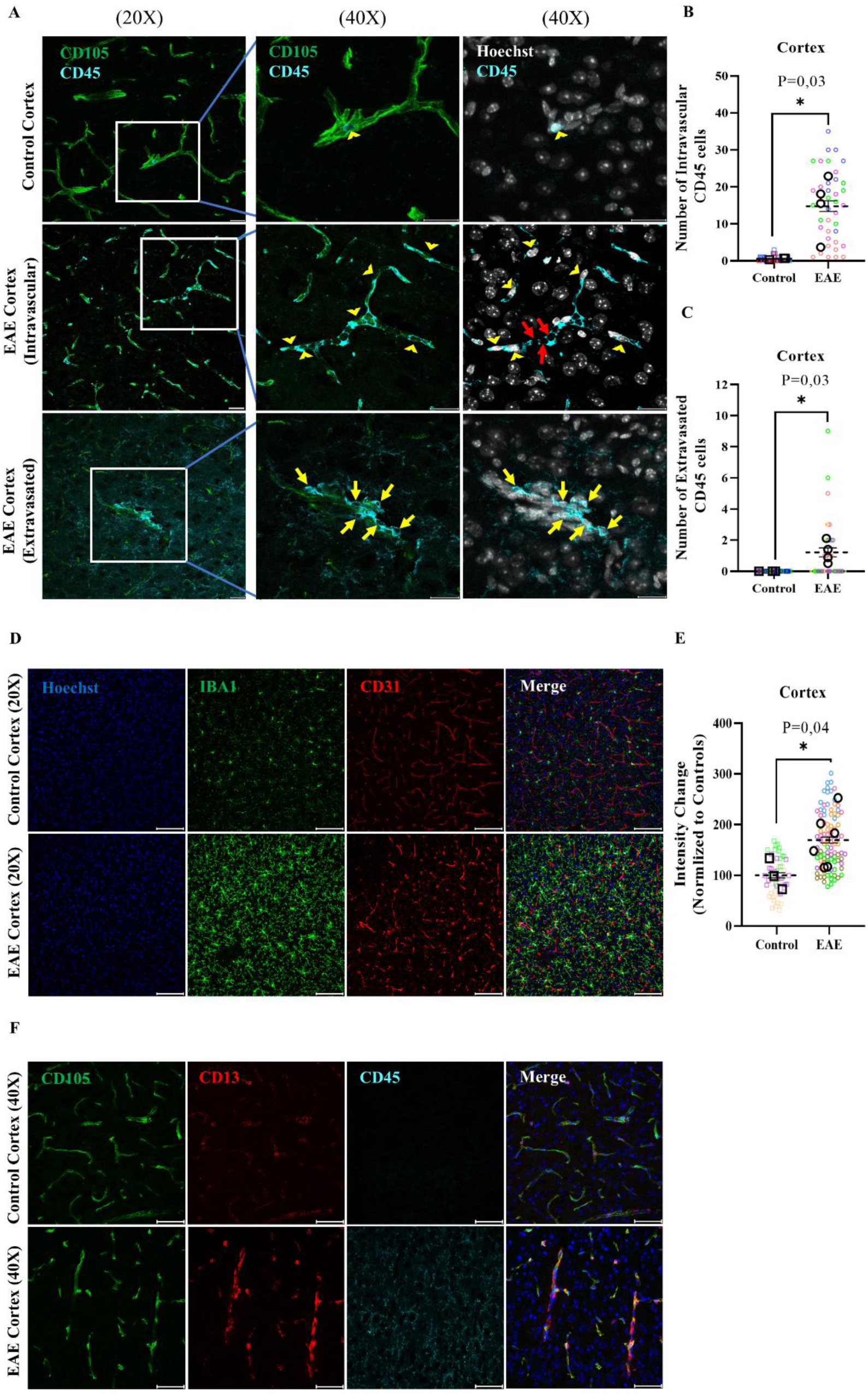
Enhanced intravascular CD45+ leukocyte stalling and microglial activation but minimal D45+ leukocytes trafficking in EAE mice due to cortical peripherally-derived inflammation. **A.** Immunofluorescent staining of CD105 (green) and CD45+ cells (cyan) reveals increased intravascular stalling (yellow arrowheads) and rare trafficking of CD45+ leukocytes (yellow arrow) through the microvasculature in the EAE mice, accompanied by increased or stalling red blood cells within the intravascular compartment, as indicated by the red arrow. Images were taken with 20X objective and magnification was performed with 40X in both the control and EAE mice. **B, C.** Quantification results indicate a significant increase in intravascular and extravasated CD45+ cells in EAE-induced mice (*p=0.03, *p=0.03, respectively). The statistical analysis was performed using a two-tailed unpaired t-test (n=3 in the control and n=4 in the EAE mice). Error bar represented mean with +/− SEM and scale bars are 20µm. **D.** Immunofluorescent staining of the cortical vasculature highlights microglial activation, marked by IBA1 (green), and platelet endothelial adhesion molecules, marked by CD31 (red), in both control and EAE-induced mice. Images were taken using 20X objective. **E.** Quantification of IBA1 intensity in the cortex reveals a significant increase in microglial activation in induced EAE mice compared to control mice (*p=0.04). Statistical analysis was performed using two-tailed unpaired t-test with n=3 in the control and n=6 in the EAE mice. Error bar represented mean with +/− SEM and scale bars are 100µm. **F.** Immunofluorescent staining of CD45+ cells (cyan), CD105 (green), CD13 (red) and Hoechst (blue) reveals that CD45 was also expressed on activated microglia under inflammatory conditions in the EAE compared to the control mice. The scale bars are representing of 50µm.

### Mild microglial activation in the EAE cortex

Until now, we have shown that low-grade inflammation affects the structure and function of the microvessels in the EAE cortex. To investigate whether there is accompanying inflammation in the parenchyma, we examined the signs of microglial activation. Indeed, there was a significant increase in Iba1+ microglial cells carrying an activated phenotype, indicating microglial activation (Fig. 5D, E; *p*=0.04). These activated microglia also expressed CD45, whereas the microglia in the control group were negative for CD45 (Fig. 5F). To assess regional differences in microglial activation, we employed tomato lectin labeling to compare the cortical and spinal cord regions, the latter being a primary site of peripheral immune cell infiltration in MOG_35-55_-induced EAE. We observed that while cortical microglial activation was significantly elevated compared to control cortex, the magnitude of activation was notably lower than that observed in the EAE spinal cord (Supplementary Fig. 6). These results demonstrate the presence of subtle, yet clear, neuroinflammation in the cortex.

### Neurodegeneration and hypoxia in the cortex of MOG35-55-induced EAE mice

So far, we have demonstrated the presence of neurovascular uncoupling and loss of cerebrovascular reactivity in association with reactive pericytes and signs of hypoxia in the cortex. As these findings are well-known causes of neurodegeneration, we next investigated the neurons and their neurites. We found a 30% reduction in NeuN+ neuronal cells in the cortex of EAE mice compared with controls, with the most pronounced loss in Layers II-IV (Fig. 6A-B; *p=0.03*). Notably, the loss of NeuN reactivity was focal in some regions, mainly in a subpial and intracortical manner (Fig. 6C). Co-staining with IgG revealed intravascular IgG deposition in some of these areas, suggesting that NeuN loss tends to occur around affected blood vessels. For comparison, we performed Neun staining of the thoracic section of ventral horn of the spinal cord. A 24% loss of NeuN+ neuronal activity was observed in EAE mice compared to control group (Supplementary Fig. 7A, B; *p=0.05*). Furthermore, anti-SMI-32 staining was used to quantify neurites, revealing a 26% decrease in neurite density in EAE mice compared to controls (Fig. 6D, E; *p=0.007*). These findings indicate ongoing neurodegeneration in the cortex concentrated around the affected blood vessels, despite the absence of overt leukocytic infiltration. Disturbances in CVR and NVC, in addition to increased leukocyte stalling, can lead to hypoxia. Therefore, we next examined the expression of HIF-1α in the cortex. We found increased HIF-1α staining around microvessels in EAE animals but not in controls (Fig. 6F).

**Figure 6.**
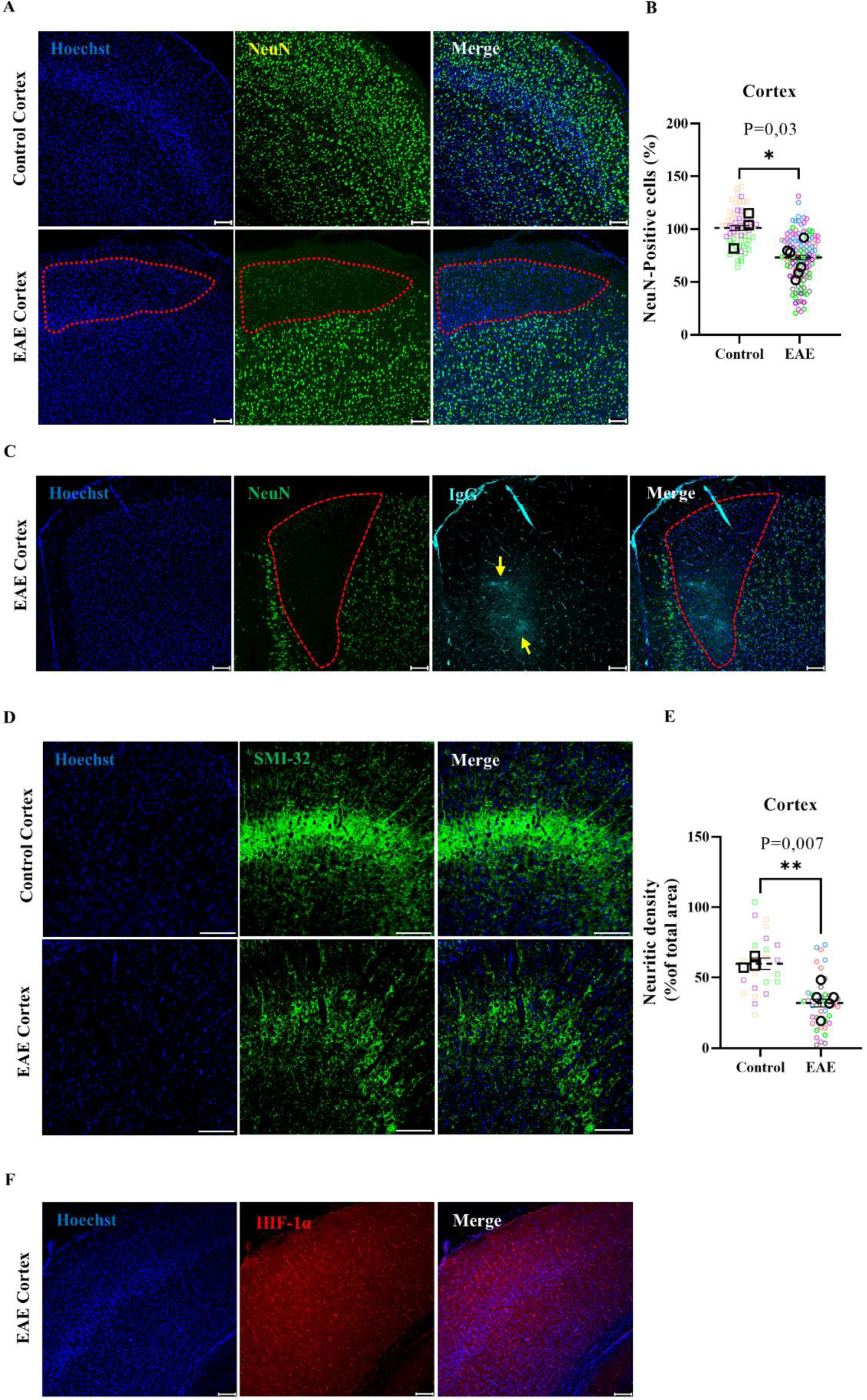
Cortical neurodegeneration in EAE mice: loss of NeuN+ neuron, decreased neurite density, and increased HIF-1α expression associated with disturbed neurovascular coupling. **A.** Confocal microscopy images indicating NeuN+ neurons (green) and Hoechst nuclear staining (blue) in the cortex of control and induced EAE mice. Images were taken using 10X objective, and neuronal counting was performed across cortical layers II-IV. The dashed red line in the EAE highlights the area of localized NeuN+ neuronal cell loss. The scale bar represents 100 µm. **B.** Quantification of NeuN+ neuron count in the cortex demonstrates a 30% decrease in the number of neurons in the cortex of induced EAE mice compared to control mice (*p = 0.03). The group size; n=3 for the control group and n=6 for the EAE group in the cortex. **C.** Immunofluorescence staining indicates a colocalization between NeuN-positive cell loss (red dashed line) and IgG deposition in the cortex. Extravasation of IgG (yellow arrowheads) is accompanied by a significant increase in luminal IgG deposition and intravascular CD45+ cells, which is associated with neuronal loss. The images were captured using 10X objectives. The scale bar representing 100 µm. **D.** SMI-32 (green) immunofluorescence staining, identifying dephosphorylated neurofilaments, reveals a reduction in neurotic density in EAE mice compared to controls. The images were captured using 20X objectives. The scale bar represents 100 µm. **E.** Quantification of Neurotic density in the cortex demonstrates a ∼ 26% decrease in the axonal neurofilament in the cortex of induced EAE compared to control mice (**p=0.007). The group size; n=3 for the control group and n=5 for the EAE group in the cortex. **F.** Double immunofluorescence staining of HIF-1α (red) and Hoechst (blue) indicate increase of HIF-1α staining around microvessels of EAE animals. The scale bars are representing of 50µm.

## Discussion

Our findings demonstrate that cortical pericytes acquire a reactive phenotype in response to CNS inflammation, alongside microglia and endothelial cells. This reactivity manifests as increased vascular coverage and compromised contractile function, evidenced by impaired cerebrovascular reactivity and neurovascular uncoupling. Importantly, we identified that pericyte dysfunction is associated with hypoxic response and neurodegeneration during neuroinflammation, suggesting a potential mechanism contributing to neurodegeneration in multiple sclerosis.

In this study, we utilized the MOG_35-55_-induced EAE model to investigate cortical pathology, an aspect that has been largely overlooked despite this being the most widely used experimental model of MS. While majority of previous studies have primarily focused on spinal cord pathology due to the presence of obvious inflammatory lesions in this region, our findings demonstrate that the cortex is significantly affected even in the absence of direct leukocyte infiltration. In our model, the administration of 135 ng pertussis toxin induced robust inflammation in the spinal cord without directly promoting leukocyte infiltration into the cortex. In a previous study, a higher dose of pertussis toxin (500 ng) combined with a double dose of MOG_35-55_ peptide was used to establish an EAE model that induced frank cortical inflammatory lesions (25). While that model demonstrated extensive BBB disruption, cortical demyelination, and neurodegeneration, our model exhibited milder cortical inflammation, characterized by a largely intact BBB and minimal leukocyte infiltration. By utilizing this approach, we were able to demonstrate that significant neurovascular dysfunction and neurodegeneration can occur even in the absence of direct leukocyte infiltration, closely mimicking the pathophysiological conditions observed in normal-appearing gray matter (NAGM) in people with MS. This is particularly relevant as cortical pathology in MS often develops without obvious inflammatory infiltrates yet contributes significantly to disease progression. Our findings therefore provide mechanistic insights into how distant inflammatory processes can trigger cortical vascular dysfunction and subsequent neurodegeneration.

The reactive pericyte phenotype we observed throughout the cortex was accompanied by increased intravascular IgG deposition, ICAM1 expression, and microglial activation, all occurring without direct leukocyte infiltration or overt BBB disruption. These findings parallel changes previously reported in NAGM in pwMS, where cortical neurodegeneration develops despite the relative scarcity of tissue-infiltrating leukocytes (26, 27). The presence of MHC class II-positive activated microglia in both active cortical lesions and normal-appearing cortex of pwMS (28, 29), further supports the clinical relevance of our model. Notably, cortical microglial activation and neurodegeneration have been shown to present a surface-in gradient, particularly in pwMS with meningeal lymphoid follicles (30). Our observation of pronounced NeuN+ neuron loss in superficial cortical layers II to IV, coupled with severe meningeal infiltration throughout the spinal cord, aligns with this pattern and suggests that inflammatory mediators likely circulate through the CSF to affect cortical structures, a mechanism particularly relevant given the anatomical characteristics of the mouse brain.

Soluble factors released into the CSF by meningeal infiltrates have been implicated in gray matter neurodegeneration in pwMS. Our in vitro findings demonstrate that H_2_O_2_, a key reactive oxygen species, directly induces upregulation of cytoskeletal and contractile molecules in pericytes, providing a mechanistic link between oxidative stress and pericyte reactivity during neuroinflammation. This observation is particularly relevant given the reported accumulation of oxidized phospholipids in cortical lesions and NAGM in pwMS (31). Additionally, inflammatory cytokines such as TNF-α, IFN-γ, and IL-1β have been associated with gray matter neurodegeneration (30), suggesting multiple pathways through which remote inflammation might influence cortical pathology.

Our study provides the first comprehensive assessment of both cerebrovascular reactivity (CVR) and neurovascular coupling (NVC) during EAE. We demonstrate that these essential vascular functions are compromised diffusely in the cortex during neuroinflammation, even without direct leukocyte infiltration. These findings are in line with several human studies. Marshall et al., using arterial spin labeling and hypercapnia induction, demonstrated that gray matter CVR is globally decreased in pwMS, with this decrease correlating directly with gray matter atrophy index (32). In their follow-up study investigating network-specific changes, they found neurovascular reactivity was impaired in four out of seven functional networks measured in pwMS in response to 5% CO₂ inhalation, with the extent of impairment correlating with both MS lesion volume and gray matter atrophy (33). Similarly, longitudinal assessments of NVC by Uzuner et al., measuring cerebral blood flow responses to visual stimulation using Doppler ultrasonography, revealed a progressive decline in NVC over a 27-week period (34). This finding was further supported by Sivakolundu et al., who reported reduced BOLD signal changes in individuals with MS compared to healthy controls, with the most pronounced deficits in patients exhibiting cognitive slowing (35).

As pericytes are principal regulators of capillary blood flow in the CNS (13, 36), our findings suggest that their dysfunction may be a primary driver of these vascular abnormalities. This aligns with our previous work showing that pericyte dysfunction causes microcirculatory deficit despite the reperfusion of large arteries after temporary middle cerebral artery occlusion (37). Furthermore, other studies have demonstrated that pericyte ablation in the brain results in neurovascular uncoupling and hypoxia, ultimately leading to neurodegeneration (38).

The role of vascular dysfunction in MS pathogenesis has gained increasing recognition in recent years (30). Geraldes et al. provided compelling evidence by demonstrating that pwMS exhibit a significantly higher prevalence of small vessel disease (39). Their detailed pathological examination revealed the presence of periarteriolar dilatation and inflammation in MS brains alongside classic demyelinating lesions. Furthermore, vascular risk factors have been identified as predictors of faster disease progression in MS, and age-related small vessel disease (SVD) has been proposed as a contributor to neurodegeneration in pwMS (40). These clinical observations indicate that cerebral small vessel disease may contribute to disease progression independent of relapse activity (PIRA), a crucial aspect of MS pathology that remains poorly understood and inadequately treated with current therapies. Our results support and expand this human data by providing experimental evidence that pericytes and cortical microvessels are immediate targets of neuroinflammation, showing both morphological and functional alterations.

The relationship between pericyte dysfunction and neurodegeneration appears to be mediated through multiple mechanisms. Our observation of increased HIF-1α expression in the cortex of EAE mice indicates tissue hypoxia, which aligns with previous studies in EAE models. For instance, upregulation of HIF-1α has been demonstrated in the spinal cord of EAE mice (41), and multispectral optoacoustic tomography studies have revealed lower oxygen saturation and compromised perfusion in the spinal cord (42). Most notably, direct PO2 measurements in the cortex and cerebellum of awake mice with MOG-induced EAE, using implanted sensors, showed decreased oxygen levels that correlated with behavioral deficits (43). The combination of compromised CVR and NVC, together with increased leukocyte stalling and elevated ICAM1 expression, creates conditions conducive to tissue hypoxia. The cortical neurodegeneration we observed, evidenced by reduction in NeuN-reactive neurons and SMI32+ neurites, likely results from these multiple convergent mechanisms, suggesting that pericyte dysfunction and neurovascular uncoupling secondary to soluble factors represent additional mechanisms contributing to cortical neurodegeneration in pwMS.

Beyond pericyte dysfunction, we identified increased leukocyte stalling associated with elevated ICAM1 expression, indicating that endothelial activation serves as an additional contributor to microvascular dysfunction and cortical hypoxia during neuroinflammation. The presence of activated microglia expressing CD45 in our model, similar to MHC class II-positive activated microglia observed in normal-appearing cortex of pwMS (27), suggests a complex interplay between vascular dysfunction and neuroinflammation.

Despite the insights provided by this study into pericyte dysfunction and neurovascular uncoupling in neuroinflammation-induced cortical neurodegeneration, several limitations should be acknowledged. First, the lack of direct examination of microvascular dynamics at the cellular level using two-photon microscopy (2-PM) limited our ability to analyze capillary flow, pericyte contractility, and leukocyte-endothelial cell interactions in real-time. Second, the absence of targeted pharmacological interventions prevented us from establishing direct causal relationships between specific pathways (such as pericyte modulation, microglial activation, or endothelial ICAM1 inhibition) and the observed vascular dysfunction, hypoxia, and neurodegeneration. Future studies addressing these limitations through 2-PM imaging and selective pharmacological interventions will be crucial for elucidating the precise mechanistic relationships between vascular dysfunction and neurodegeneration in MS.

In conclusion, our study demonstrates that cortical pericytes undergo significant functional and morphological changes during neuroinflammation, even in the absence of direct leukocyte infiltration. The increased pericyte coverage, coupled with compromised cerebrovascular reactivity and neurovascular uncoupling, represents a novel mechanism contributing to cortical pathology in neuroinflammatory conditions. These vascular alterations, accompanied by increased leukocyte stalling and microglial activation, ultimately lead to tissue hypoxia and neurodegeneration. The observation that these changes occur in normally appearing cortex, driven by remote inflammatory processes, provides new insights into the mechanisms of cortical damage in multiple sclerosis. These findings suggest that targeting pericyte dysfunction and vascular abnormalities might represent a promising therapeutic strategy to prevent cortical neurodegeneration in people with MS, particularly during the progressive phase of the disease when conventional immunomodulatory treatments show limited efficacy. Future studies employing two-photon microscopy and targeted pharmacological interventions will be crucial to further elucidate the mechanistic relationships between pericyte dysfunction, vascular compromise, and neurodegeneration in multiple sclerosis.

## Data availability

The data such as MATLAB codes that support the findings of this study are available from the corresponding author, upon reasonable request.

## Acknowledgements

We extend our sincere gratitude to TUBITAK 1001 (grant No: 121S588) and TÜBİTAK 2236 (grant No: 119C018) for providing the financial support that made this research possible. Additionally, we acknowledge the European Federation of Immunology Society (EFIS) for funding Mohammadreza Yousefi’s two-month research visit to Ludwig-Maximilian University (LMU), Munich, Germany.

We are also indebted to the KOÇ University Research Center for Translational Medicine (KUTTAM) for facilitating the study through their services and infrastructure.

## Supplementary material

Supplementary material is available online.

## Supplementary Figure legends

**Supplementary Figure 1.**
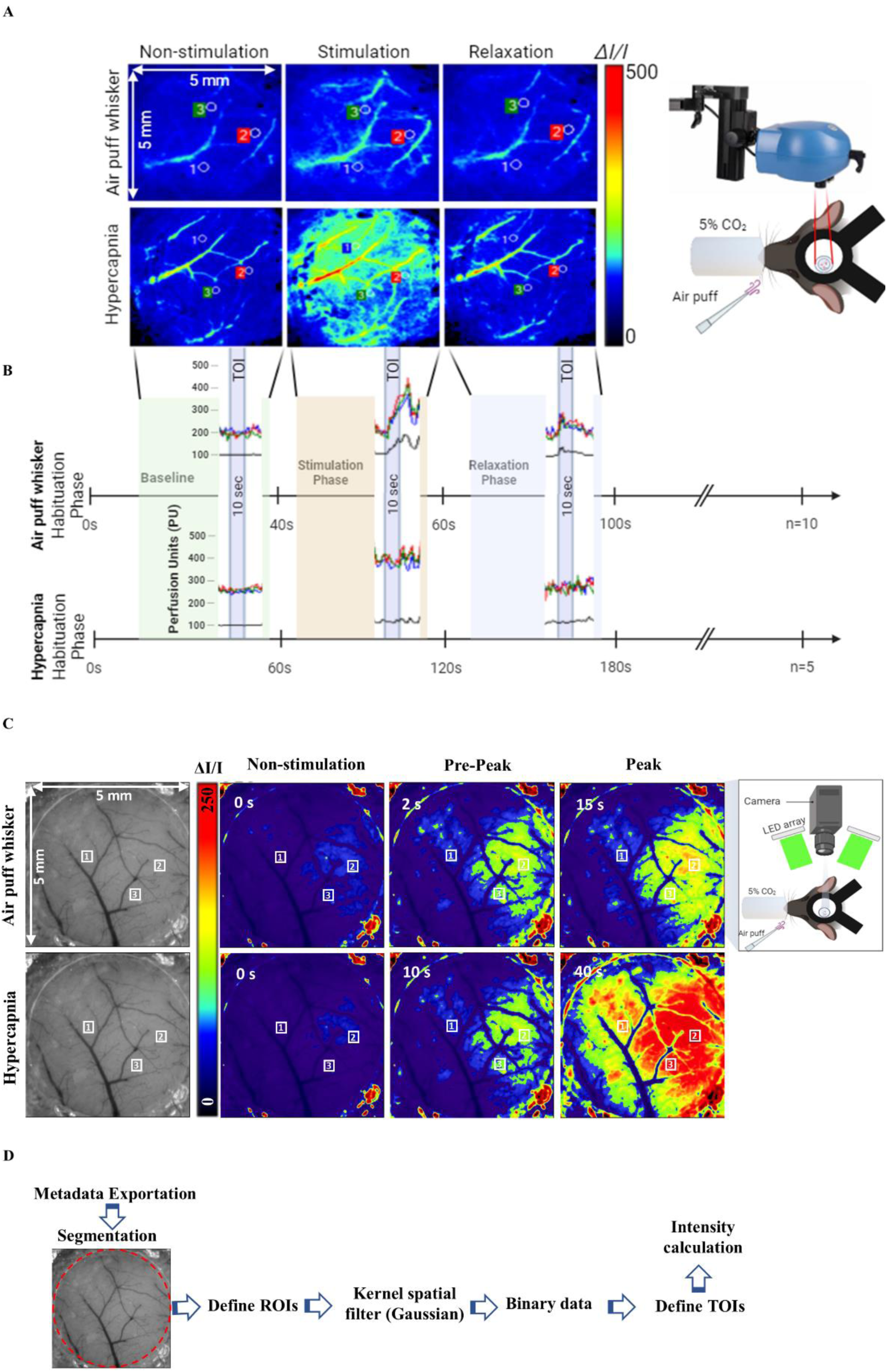
Experimental setup and data analysis workflow for air puff whisker pad stimulation and hypercapnic conditions. **A.** Representative LSCI heatmaps demonstrate relative fluorescence intensity change of perfusion (ΔI/I) in response to air puff whisker pad stimulation (top row) and 5% hypercapnic (bottom row) condition. Three different 10×10-pixel ROIs beyond the rim or slightly farther from large vessels (0.3–0.8 mm away from arterioles) were manually selected. The obtained images represent three distinct phases: non-stimulation, stimulation, and relaxation. The right panel indicates the experimental setup, comprising of an air puff administration system combined with 5% CO_2_ revealing for hypercapnic conditions. The color scale represents the intensity changes which were automatically obtained from Pimsoft software **B.** Time course analysis of the blood perfusion changes during the baseline, stimulation, and relaxation timelines. The top timeline represents the air puff and bottom timeline corresponds to the hypercapnic conditions. Vertical dashed lines imply the transition between different phases, and the colored area represents the stimulation phases. The data resulted from averaging the interaction between 10 sec TOI and ROI across multiple trials (n=10 for air puff; n=5 for hypercapnia). **C.** IOSI images and heatmaps exhibit relative fluorescence intensity change of perfusion (ΔI/I) in response to air puff whisker pad stimulation (top row) and 5% hypercapnic (bottom row) condition. All other procedures outlined in section A were operated in the same manner here. The color scale represents the intensity changes manually obtained by using Image J software. **D.** Data analysis workflow for image analysis of metadata exportation, segmentation to define ROIs, application of a Gaussian spatial filter, conversion to binary data, and intensity calculation by defining TOI intervals were utilized by customized MATLAB code.

**Supplementary Figure 2.**
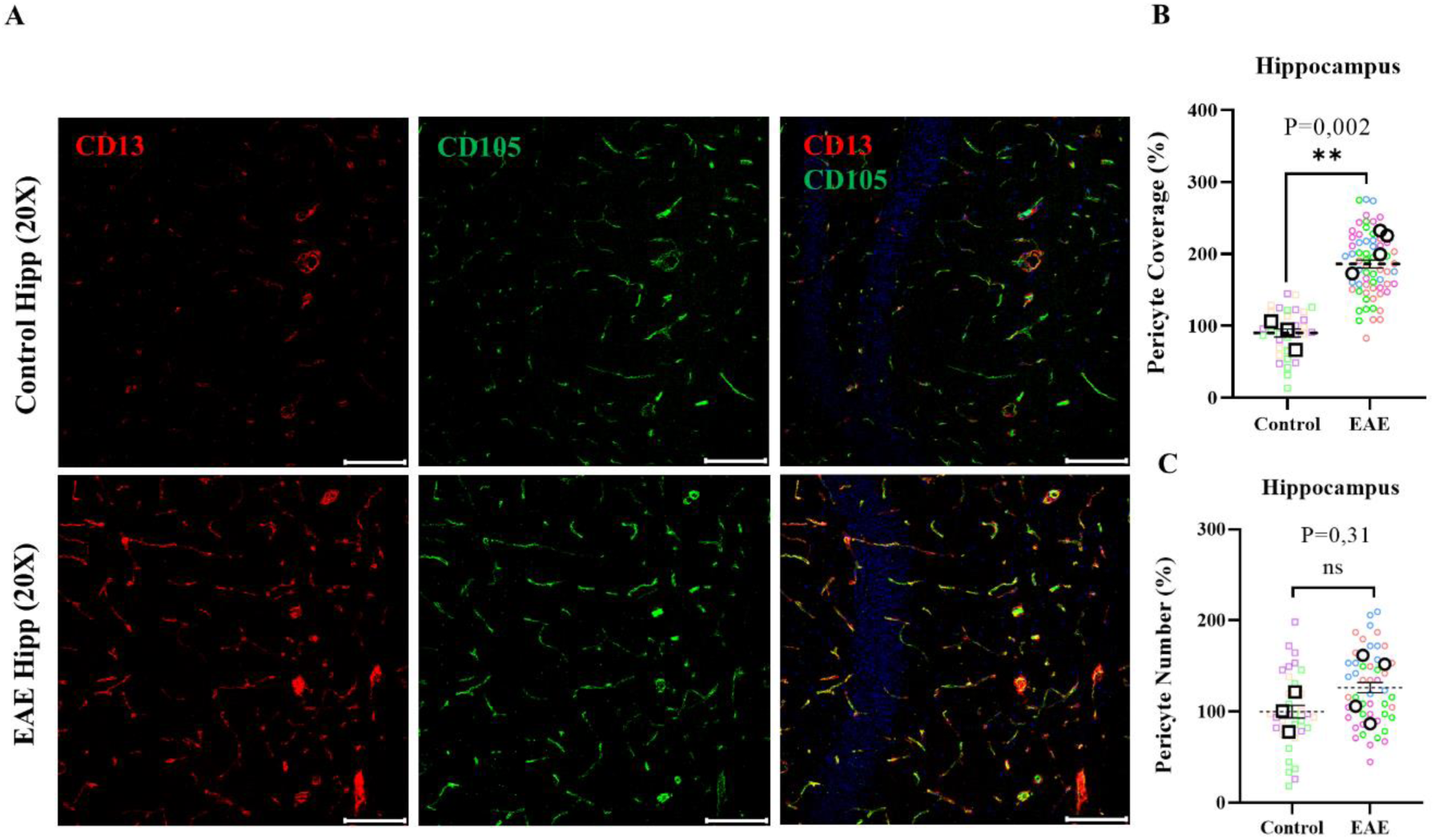
Pericyte coverage is increased in response to inflammation around hippocampal microvessels during EAE. **A.** Immunofluorescent staining reveals pericyte coverage (CD13, red) and endothelial proliferating cells marked by CD105 (vasculature, green) in the cortex of control and induced EAE mice. Representative confocal images were taken using 20X objectives in both control and EAE mice. **B.** Quantification of vascular pericyte coverage reveals a %118 increase in the hippocampus of induced EAE mice compared to control mice (**p=0.002). **C.** Quantification of the number of pericyte cell bodies on the microvasculature demonstrates no significant difference in the hippocampus of induced EAE mice compared to control mice (ns p=0.31). Statistical analysis was performed using a two-tailed unpaired t-test, with n=3 in the control group and n=4 in the EAE group. Error bars represent the median with interquartile range. Scale bars in all images represent 50µm.

**Supplementary Figure 3.**
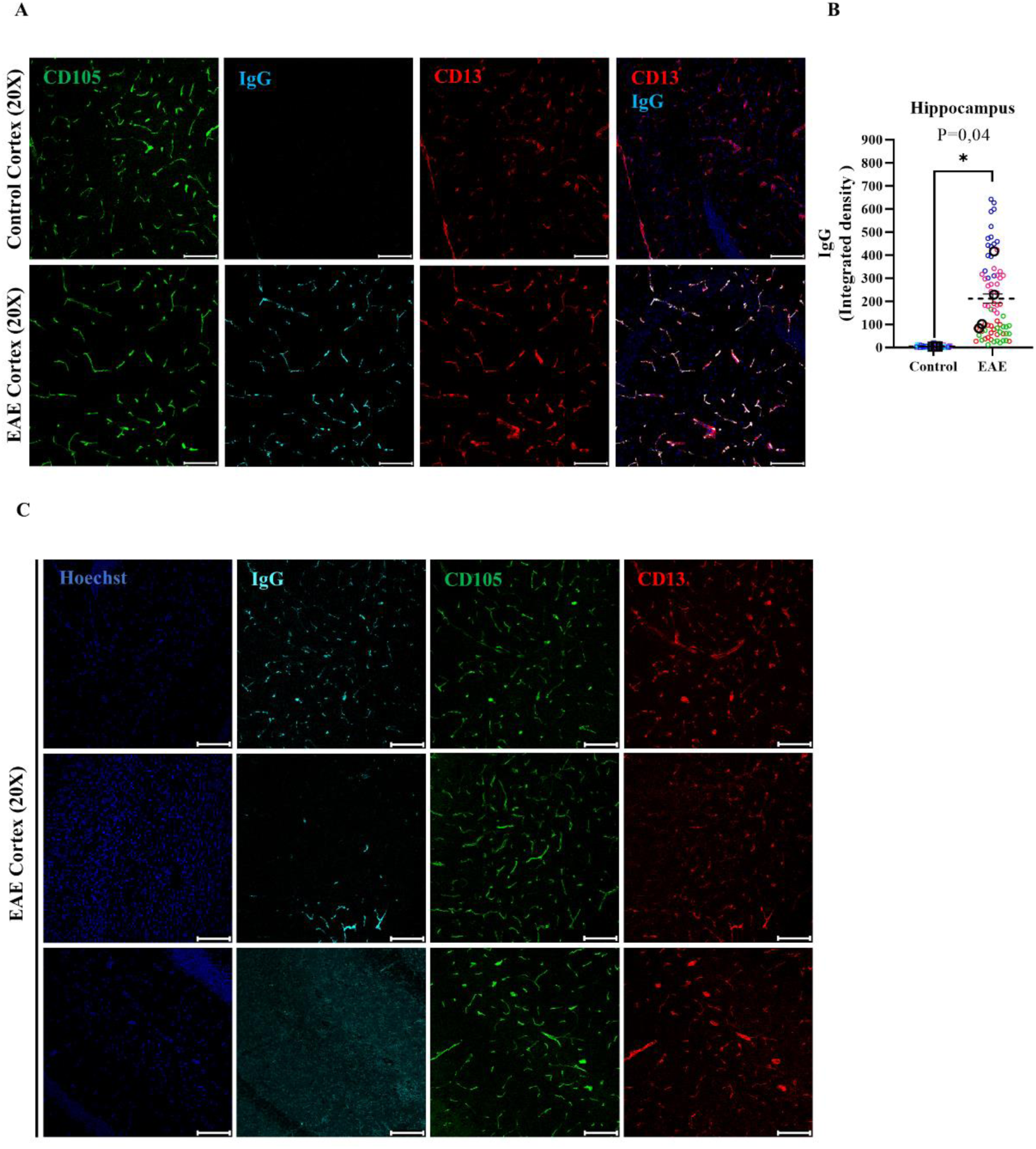
Intraluminal IgG depositions are increased in Hippocampal microvessels during EAE. **A.** Confocal microscopy images show plasma-derived IgG stained (cyan), along with CD105-stained vessels (green) and CD13-stained pericytes (red), in the hippocampus of control and induced EAE mice. Images were taken by 20X objectives. **B.** Quantification of plasma-derived IgG in the hippocampus reveals a substantial increase in luminal IgG deposition in induced EAE mice compared to control mice (*p= 0.04). **C.** with meticulous observation, plasma-derived IgG shows deposition in both focal and diffuse area within the cortex and hippocampus, specifically in the luminal region of the microvasculature. Statistical analysis was performed using a two-tailed unpaired t-test, n=3 in the control group and n=4 in the EAE group. Error bar represents median with interquartile range. Scale bars are 50µm.

**Supplementary Figure 4.**
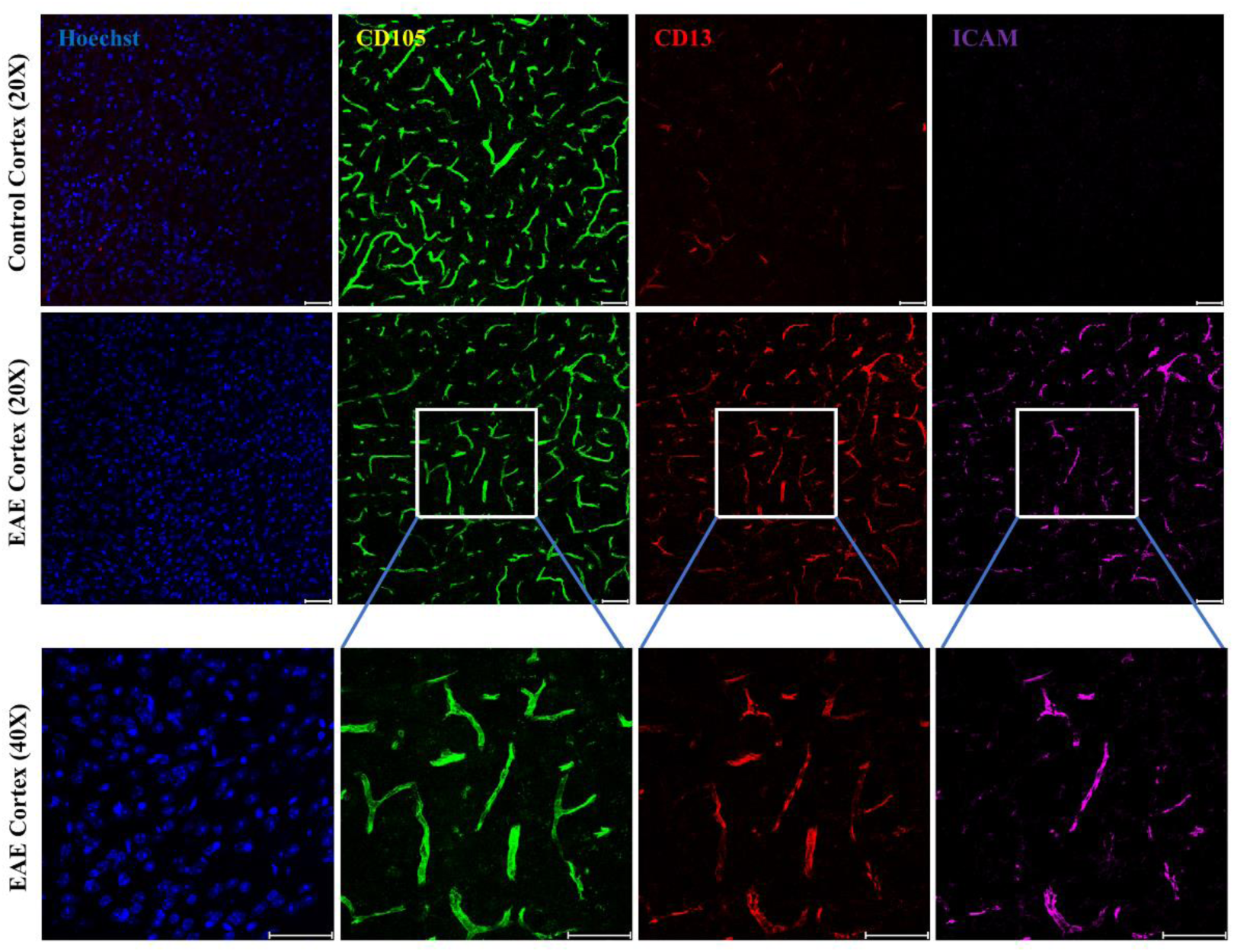
Endothelial Intracellular Adhesion Molecule 1 expression was markedly increased on the cortical microvessels in the EAE group compared to control mice. Captured immunofluorescent images were indicated staining of CD105 (green), CD13 (red), CD45+ cells (cyan) and Hoechst nuclear staining (blue) in the cortex of induced EAE mice. 20X magnification for control and EAE groups and 40X for the higher magnification were used in EAE mice. Scale bars representing 20µm.

**Supplementary Figure 5.**
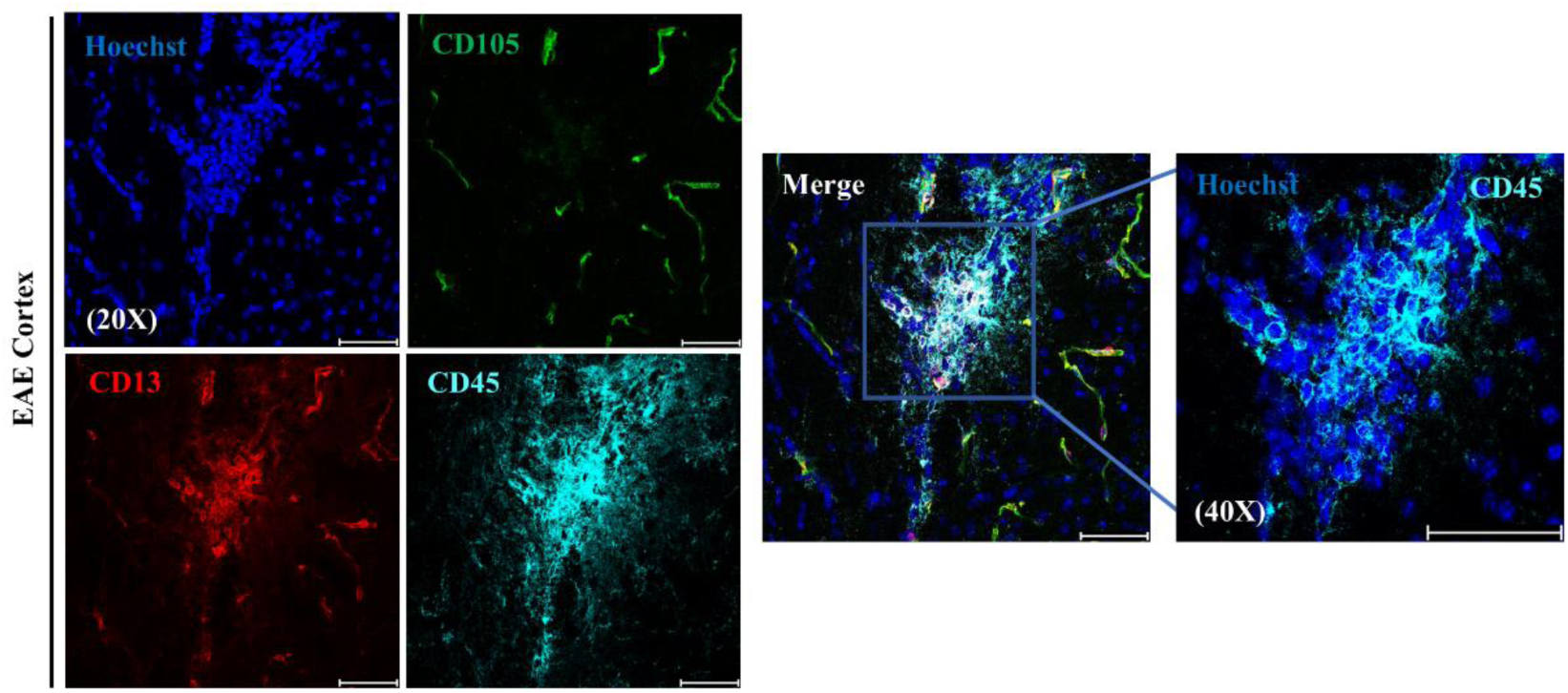
Immunofluorescent detection of extravascular CD45+ leukocytes and infiltration in the cortex of EAE mice. Representative confocal images show immunofluorescent staining of CD105 (green), CD13 (red), CD45+ cells (cyan) and Hoechst nuclear staining (blue) in the cortex of induced EAE mice. The images were captured using 20X and 40X objectives, with scale bars representing 50µm.

**Supplementary Figure 6.**
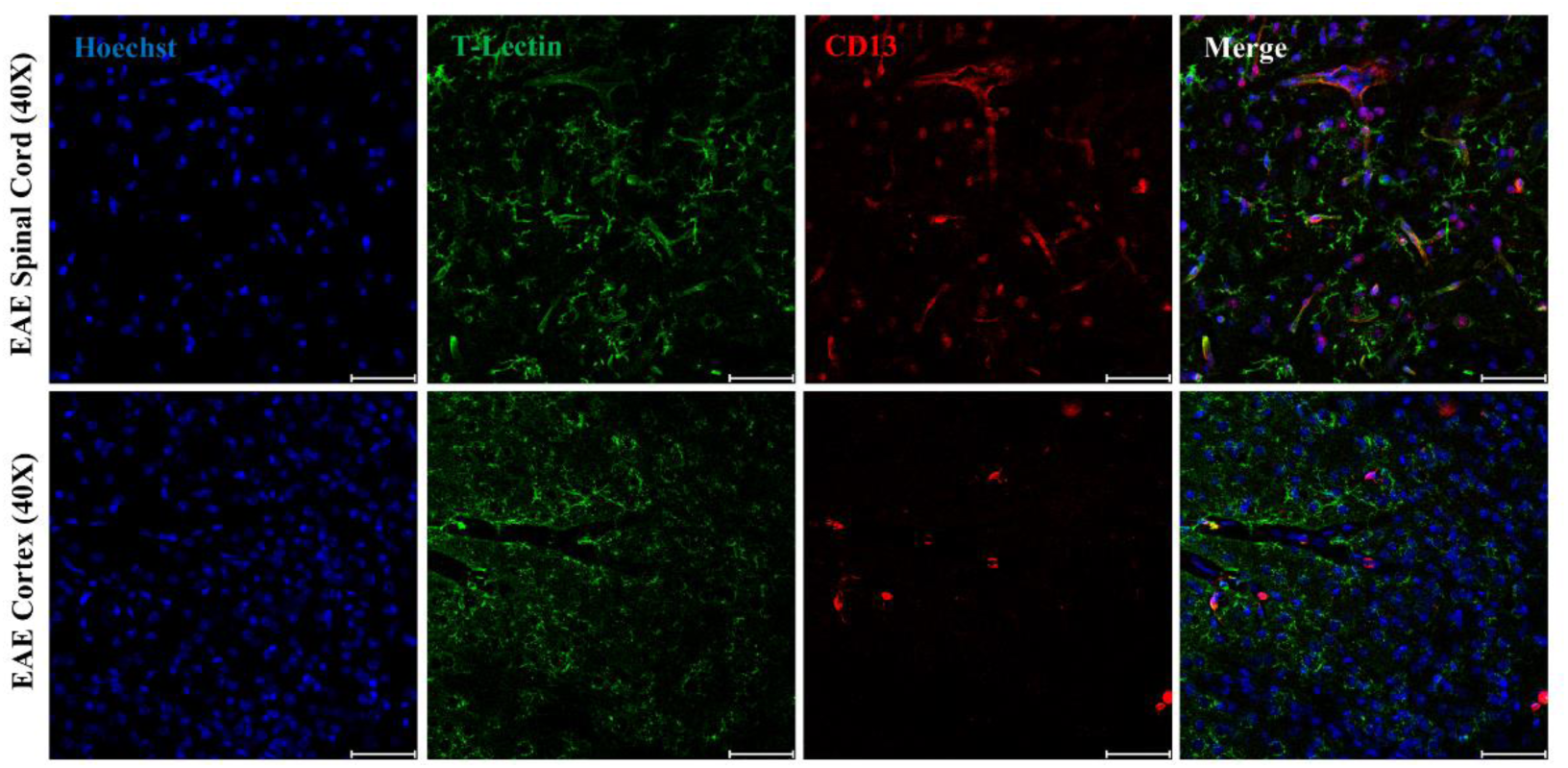
Representative confocal images of the spinal cord (top row) and cortex (bottom row) indicate immunoflourecent staining of activated microglial with tomato lectin (green), CD13 (red), and Hoechst nuclear staining (blue) in both spinal cord and cortex of the induced EAE mice. The images were captured using 40X objectives, with scale bars representing 50µm.

**Supplementary Figure 7.**
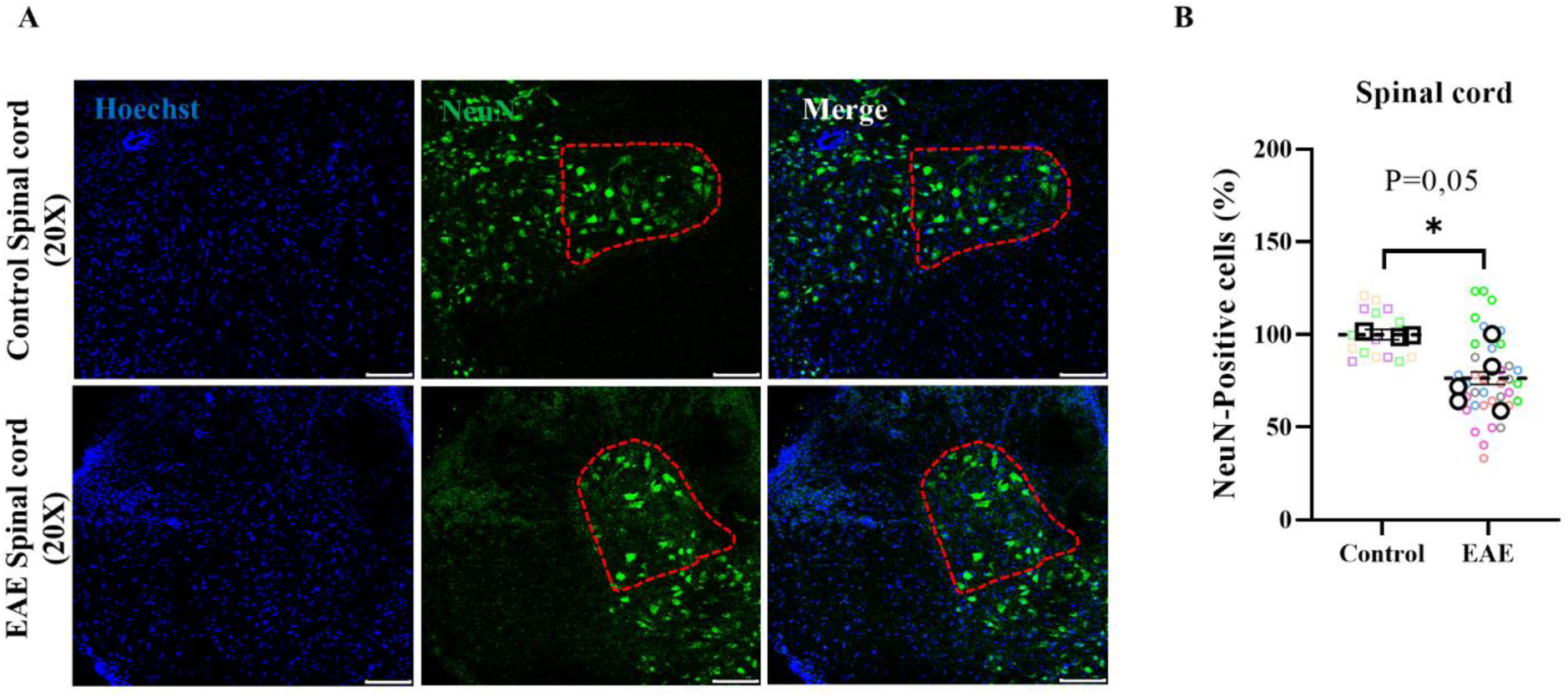
Validation of cortical NeuN+ cell loss in relation to spinal cord contribution to neuronal degeneration in the EAE model. **A.** neurons (green) and Hoechst nuclear staining (blue) in the cortex of control and induced EAE mice. The images were taken from the thoracic section of the spinal cord ventral horn using a 10X objective, and neuronal counting was performed across cortical layers II-IV. The ventral horn, outlined by the red dashed line in the EAE group, marks the area of localized NeuN+ neuronal cell loss. The scale bar represents 100 µm. **B.** Quantification of NeuN+ neuron count in the spinal cord ventral horn demonstrates a 24% decrease in the number of neurons in the spinal cord ventral horn of induced EAE compared to control mice (*p = 0.05). The group size; n=3 for the control group and n=5 for the EAE group in the cortex.

**Table 1.**
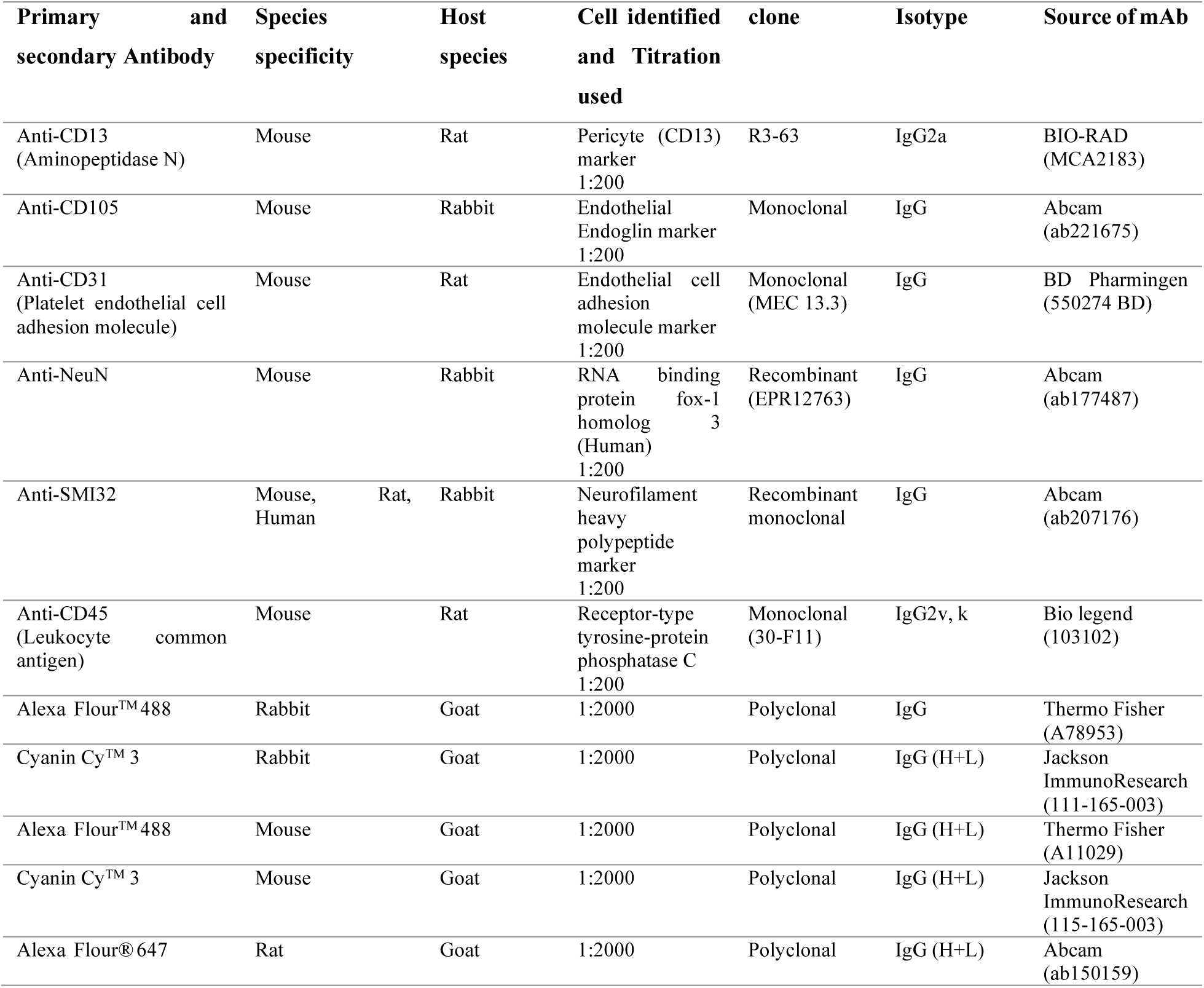
List of primary and secondary antibodies used for immunofluorescent staining.

**Table 2.**
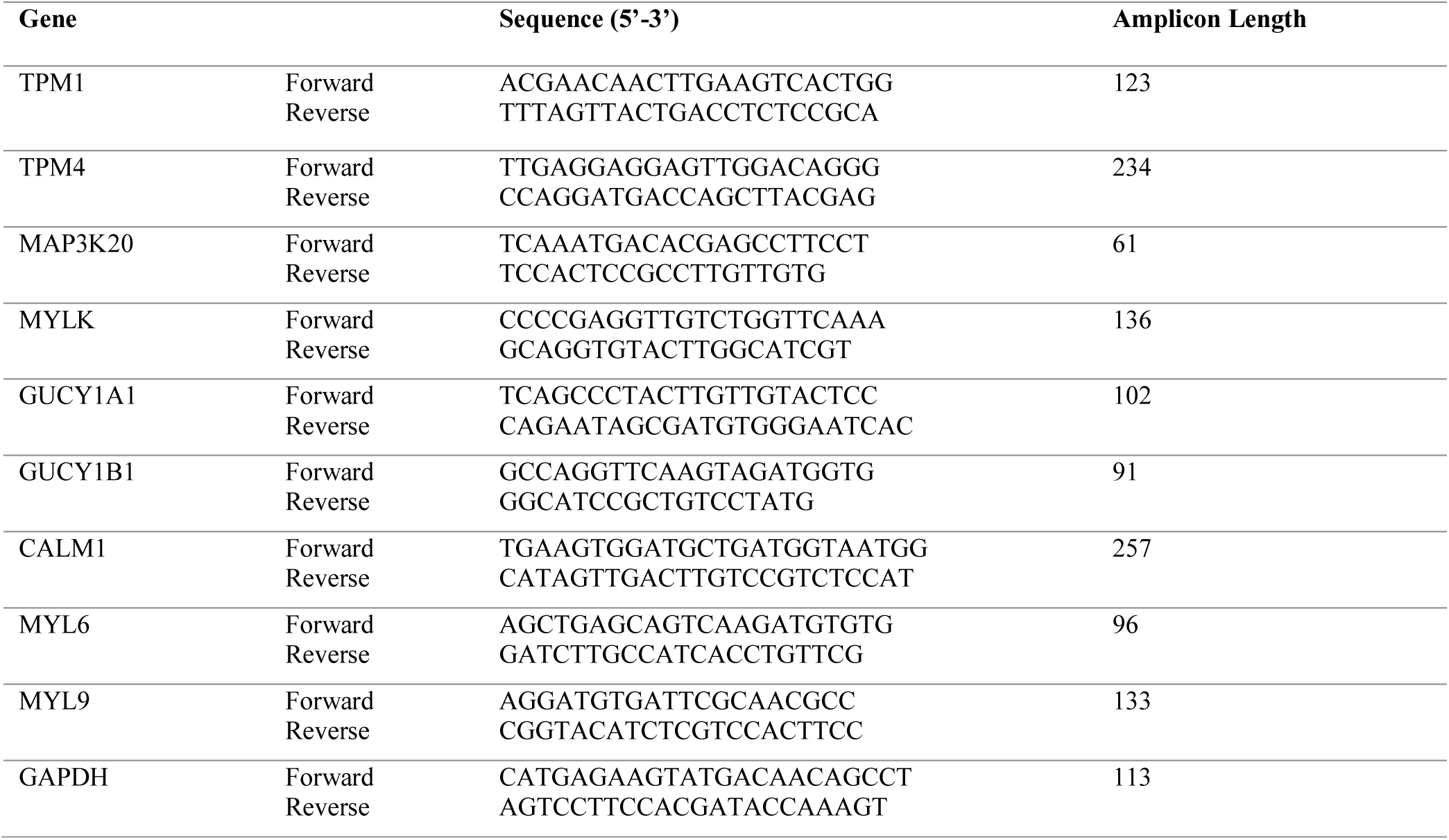
List of primers used for RT-PCR.

