## Supplementary Methods for "Reactive Pericytes Lead to Microvascular Dysfunction and Cortical Neurodegeneration During Experimental Autoimmune Encephalomyelitis"

**Custom-made plug-in glass coverslip.**

Following the best possible optimized results, a few days after the surgery, the anti-inflammatory medication efficacy seems to have decreased. This decrease may have caused edema to develop, which might have put pressure on the brain against the edge of the teared skull section and eventually caused bleeding within the cranial window. To address this limitation and mimic a skull thickness of ~300µm, a custom-made plug-in glass coverslip was adopted. This required assembling of a three separate, roughly 100 µm-thick pieces of 3mm cover glass. UV-curved optical glue (NOA61) was used to carefully attach these parts to 5mm slides. Our prepared plug-in glass coverslip, upon completion of the cleaning and sterilization, were meticulously affixed to the window.

**Custom-made head plate.**

Since the required apparatus for the awake experiment, such as the head plate, might present certain drawbacks and not be appropriate, it was specially designed in a certain size with computer-aided design software and produced with 3D printing techniques (see Supplementary Figure). The 123Desing tool was used to design the component to ensure that it fulfilled the necessary dimensions. After that, it was formatted in STL to make it compatible. By using the Ultimaker CURA-Slicer software, the resultant STL file was then converted to G-code, transforming it into a readable format that is appropriate for the 3D printer. The printer that was used, specially designed for this particular production, has an accuracy of 25 microns on the X, Y, and Z axes. A 3D model was created using the Fused Deposition Modeling (FDM) process. The temperature at which the polymeric material was produced was 250 degrees for the extruder and 60 degrees for the hotbed. The produced parts were cooled to room temperature after printing and then sterilized with ethyl alcohol and UV. Finally, the head plates were packaged and stored to be applied on the animals.

**Custom-made head plate restraint.**

Taking the head plate design as a foundational framework, the restraint device was designed with the express purpose of immobilizing the awake mouse during the imaging. For this purpose, the parts planned to be used for the experiment were specially designed through the 3D design software 123Desing to fulfill multiple functions suitable for the microscope and working area (see Supplementary Figure). the holding system, which is designed and manufactured in accordance with the head plates, can be adjusted according to the position of the animal, the desired experiment shape and height of the microscope. Furthermore, the presence of perforations on the holder floor, facilitate development to accommodate different experiments envisioned for the future.

**Custom-made air puff whisker pad.**

As specialized air puff system was created and used in this work to provide mice’s whisker area with regulated stimuli. The apparatus regulates a solenoid valve (air puff motor) that manages the flow of compressed gas (CO2) using an Arduino microcontroller. To guarantee consistency in stimulus delivery, the Arduino was configured to produce the exact timing sequencing for the air puffs, which are supplied at predetermined intervals. The valve’s actuation was controlled by a motor driver, which allowed for frequency, duration and count adjustment. This configuration appropriate for behavioral and neurophysiological research since it guarantees precise, reproducible whisker stimulation.to reduce the possibility of vibration and noise when the solenoid valve is operating, it was applied remotely using a 6 mm thick PTEE (Teflon) pipe that is appropriate for 1 bar. The device was designed to sustain strong and consistent activation of the sensory in the whisker pad region while offering a moderate and non-invasive stimulus to prevent pain.

**Custom-made IOSI.**

A non-invasive imaging method called IOSI that uses variation in light reflectance rom the brain tissue to gauge brain activity. These alterations are linked to physiological functions, specifically oxy-deoxygenation, blood flow, and neuronal activity.by tracking variation in oxy-deoxy hemoglobin levels, we used an IOSI to identify particular region of the brain activation. A specially made LED array with wavelength of 530 and 630 nm is used in the system. These wavelengths-530 nm for detecting changes in oxyhemoglobin and 630 nm for deoxyhemoglobin were selected because of their sensitivity to the oxygenation state of hemoglobulin. The brain surface’s reflected light was recorded, and the difference in absorption at various wavelength gave a precise indication of how local blood oxygenation changed in response to neuronal activity. We obtained a high signal-to-noise ratio (SNR) by employing dual-wavelength illumination, which enables accurate function area detection with strong sensitivity to homodynamic changes. Accurate mapping of the brain regions linked to whisker stimulation was made possible by this technique, which allowed for real-time imaging of the neurovascular response to sensory stimulation.

**CD13-Positive Pericyte imaging and coverage analysis.**

The images were captured at 20X objective with a 2-micron-apart Z-stack of 10–15 microns thick maximum projection by a Leica DMI8 SP8 confocal microscope. In each animal, 3-5 randomly selected areas from the cortex and 3-6 randomly selected areas from the hippocampal region (581 x 581 µm) were analyzed in each of three separate, non-adjacent sections (60 microns apart) by a blinded investigator. To excite and detect the emission of Alexa Flour 488, a 488 nm laser and a 500–50 nm detector was used, while to excite and detect the AF-647, a 633 nm laser and a 660–720 nm detector was used. Then the images were completely analyzed using NIH ImageJ. For analysis of the PC coverage, every captured image was transformed into the maximum projection image using LASX software. Subsequently, the preserved Z-stack images were individually analyzed by converting them to 8-bit format, applying a Gaussian filter to enhance signal intensity and using triangle thresholding to quantified integrated density. To determine the coverage percentage, the integrated density of CD105 signals was divided by the integrated density of CD13 signals, considering only images with a resolution bigger than 50 pixels to exclude background noise. 4-6 animals in experimental group and 3 animals in control groups were analyzed in cortex and hippocampus regions, respectively.

**CD13-Positive Pericyte imaging and Count analysis.**

The conditions described in the coverage analysis were applied for taking and preparation of the images. PCs were identified by the CD13-positive staining cells that overlapped with a nucleus of cell body that was present on the CD105-positive cell signal. To prevent repeating the projections of the counted PCs, the counting was done by repetitively checking through several pages by a two blinded investigator in the single cell analyzed tool using NIH ImageJ. In order to compare the number of PCs in each image between two groups, the number of counted PCs in each image was normalized into the CD105 vessel percentage area. 4-5 animals in experimental group and 3 animals in control groups were analyzed in cortex and hippocampus regions, respectively. CD-13 positive cells counting were done by a two blinded investigator.

**Imaging and analysis of Neurotic density with SMI-32 positive cells.**

Images were captured using a Leica DMI8 SP8 confocal microscope at a 10X objective, with a 2-micron-apart between z-stack images, each having a maximum projection thickness of 10-15 microns. In each animal, two randomly selected ROIs of SMI-32 positive area (980 x 980 µm) from both of retro splenial cortex (RSP) and posterior parietal association (PPA) areas of the cortex were analyzed in each of three separate, non-adjacent sections (60 microns apart). To excite and detect the emission of Alexa Flour 488, a 488 nm laser and a 500–50 nm detector was used. The assessment of SMI-32 neurotic density was quantified by exporting 3D images, applying threshold processing, and calculating the percentage of SMI-32 positive regions within the brain by a blinded investigator using the ImageJ area measurement tool. 5 animals in experimental group and 3 animals in control groups were analyzed in cortex.

**Imaging and analysis of Neuronal Nuclei Count with NeuN+ cells.**

The images were captured at 10X and 20X (cortex and spinal cord, respectively) objective with a 2-micron-apart Z-stack of 10–15 microns thick maximum projection by a Leica DMI8 SP8 confocal microscope. NeuN-stained channel images were analyzed with ImageJ cell counter tool to quantify NeuN-positive neuronal nuclei. Since the number of neurons in the cortex layers of different brain areas and the morphology of the ventral horn in each section of the spinal cord varies, the three adjacent brain and spinal cord section (60 micron apart) were used and counted only the layers of the cortex from II-IV and ventral horn of the thoracic section of the spinal cord. In each animal, 3–4 randomly selected cortex fields (1162x1162 µm) and from each field 1-3 distinct ROIs (340 x 340 µm) were selected and analyzed by a blinded investigator using the ImageJ cell counter tool. In addition, randomly 5-6 ventral horn fields containing NeuN-positive cells were selected and analyzed. To excite and detect the emission of Alexa Flour 488, a 488 nm laser and a 500–50 nm detector was used. 5-6 animals in experimental group and 3 animals in control groups were analyzed in cortex and spinal cord regions, respectively.

**Imaging and analysis of luminal IgG deposition.**

The images were captured at 20X objective with a 2-micron-apart Z-stack of 10–15 microns thick maximum projection by a Leica DMI8 SP8 confocal microscope. In each animal, 2-4 randomly selected fields from the cortex and 4 randomly selected fields (581 x 581 µm) from the hippocampal region were analyzed in each of three separate, non-adjacent sections (60 microns apart). Triangular thresholding was used to measure the signal intensity to determine the integrated density of the luminal IgG in the microvasculature by a blinded investigator using the ImageJ. To excite and detect the emission of CY3, a 561 nm laser and a 550–600 nm detector was used. 4 animals in experimental group and 3 animals in control groups were analyzed in cortex and hippocampus regions, respectively.

**Imaging and analysis of CD45+ cells.**

Images were captured using a Leica DMI8 SP8 confocal microscope at a 40X objective, with a 2-micron-apart between z-stack images, each having a maximum projection thickness of 10-15 microns. 4 animals in experimental group and 3 animals in control groups were selected and 3-4 randomly selected field (290 x 290 µm) from the cortex were analyzed in each of three separate, non-adjacent sections (60 microns apart). Using selected 40X images, CD45-positive leukocytes were identified by the CD45 specific marker staining that overlapped with a nucleus of cell body that was present inside and outside the CD105-positive cell signal. To prevent repeating the projections of the counted CD45-positive leukocytes, the counting was done by repetitively checking through several pages in the single cell analyzed tool using NIH ImageJ. CD45-positive cells counting were done by a two blinded investigator. To excite and detect the emission of Alexa Flour 488, a 488 nm laser and a 500–50 nm detector was used.
